# “Single-nucleus RNA-seq2 reveals a functional crosstalk between liver zonation and ploidy”

**DOI:** 10.1101/2020.07.11.193458

**Authors:** M. L. Richter, I.K. Deligiannis, A. Danese, E. Lleshi, P. Coupland, C.A. Vallejos, M. Colome-Tatche, C.P. Martinez-Jimenez

## Abstract

Single-cell RNA-seq reveals the role of pathogenic cell populations in development and progression of chronic diseases. In order to expand our knowledge on cellular heterogeneity we have developed a single-nucleus RNA-seq2 method that allows deep characterization of nuclei isolated from frozen archived tissues. We have used this approach to characterize the transcriptional profile of individual hepatocytes with different levels of ploidy, and have discovered that gene expression in tetraploid mononucleated hepatocytes is conditioned by their position within the hepatic lobe. Our work has revealed a remarkable crosstalk between gene dosage and spatial distribution of hepatocytes.

## Background

The liver performs a wide variety of physiological functions, including metabolism of xenobiotics [1–5] and the regulation of energy homeostasis [6, 7], among others. These functions are mostly performed by hepatocytes which constitute 70% of the parenchymal cells. Liver non-parenchymal cells (NPCs), namely, liver endothelial cells, biliary cells, kupffer cells, hepatic stellate cells and immune cell populations constitute the remaining 30% of cell types. All these cells are organized in repetitive structures named liver lobules [8, 9]. The analysis of individual cells by single-cell genomics is changing our understanding of liver homeostasis and pathogenic conditions by taking into account their spatial distribution [10–16]. However, single-cell isolations from the liver require harsh enzymatic or mechanical dissociation protocols that perturb the mRNA levels [17]. In particular, the two-step collagenase perfusion generally used to isolate hepatocytes from human livers leads to the downregulation of liver-specific transcription factors such as *Hnf4a, Cebpa, Hnf1a,* and *Foxa3* [18], as well as their downstream target genes such as *Cyp2c9, Cyp2e1, Cyp2B6, Cyp2D6, Cyp3A5* and *Cyp3A4* [17, 19].

Single-nucleus RNA-seq (snRNA-seq) has emerged as a complementary approach to study complex tissues at a single-cell level [20, 21], including brain [21–27], lung [28], kidney [29–32] and heart [33, 34] in mouse and human frozen samples [35, 36]. However, there are no snRNA-seq methods tailored for frozen liver tissues. The nuclear transcriptome of individual cells has shown a high correlation to the cytoplasmic RNA [23, 37] indicating that single-nucleus RNA-seq is a powerful tool to study tissues from which intact and fresh cells are difficult to obtain. Here, we have developed a robust single-nucleus RNA-seq2 (snRNA-seq2) approach that relies on an efficient lysis of the nuclear membrane. Our approach permits the unbiased characterization of all major cell types present in the liver from frozen archived samples with high resolution.

With snRNA-seq2, we explore at the single-cell level a defining feature of hepatocytes, ploidy [38–43]. At birth all hepatocytes are diploid, with a single nucleus containing two copies of each chromosome. During development, polyploidization gradually increases, leading to hepatocytes with several levels of ploidy. Hepatocyte ploidy depends on the DNA content of each nucleus (e.g. diploid, tetraploid, etc.) and the number of nuclei per hepatocyte (e.g. mono- or bi-nucleated) [40]. Here we present a novel analysis of diploid (2n) and tetraploid (4n) nuclei from the mouse liver and demonstrate that ploidy is an additional source of hepatocyte heterogeneity, connecting gene dosage and liver zonation.

## Results

### snRNA-seq2 allows deep and robust characterization of single nuclei isolated from frozen livers

In order to explore archived samples associated with healthy and disease conditions, we have developed a robust methodology that combines transcriptomics and efficient low-volume reactions in single nuclei isolated from frozen livers.

Purified diploid (2n) and tetraploid (4n) nuclei were FACS sorted in an unbiased fashion according to their genome content, followed by a modified version of Smart-seq chemistry using liquid handling robots for volume miniaturization (Fig. 1A, see Methods). This approach allows the detection of over 400,000 reads per nucleus, leading to an average detection of more than 11,000 transcripts (4,000 genes) per nucleus (Supp. Fig. S1A-B). As expected for the nuclear transcriptome, the percentage of reads mapping to intronic regions were more than 68%, and the ribosomal reads were 1.2% (Fig. 1B). To show the robustness and reproducibility of this method, livers from four biological replicates harvested from young mice (3 months old) were analyzed, including two technical replicates. The technical replicates consisted of nuclei isolated from the same frozen liver on different days (Supp. Fig. S1A and Methods). Additionally, ERCC RNA spike-in mix was used to address technical noise and account for plate effects (including library preparation and sequencing) [44, 45]. ERCC normalization was used to make counts comparable across cells and minimize results purely driven by ploidy [46–49] (Methods; Supp. Fig. S1A). SnRNA-seq2 detected on average more than 4000 genes, which is a high number of genes detected per nucleus compared with other single-cell approaches in which intact cells are isolated from livers (Fig. 1C and Supp. Fig. S1B). Furthermore, our approach showed a Pearson correlation of 0.62 between gene expression in the nuclei using our snRNA-seq2 and scRNA-seq from intact cells isolated from fresh livers (Supp. Fig. S1C). This correlation shows that single-nucleus RNA-seq2 is a robust and reliable approach to study transcriptional profiles of individual cells from archived frozen tissues [20, 23, 25, 36, 50–52].

**Figure 1.**
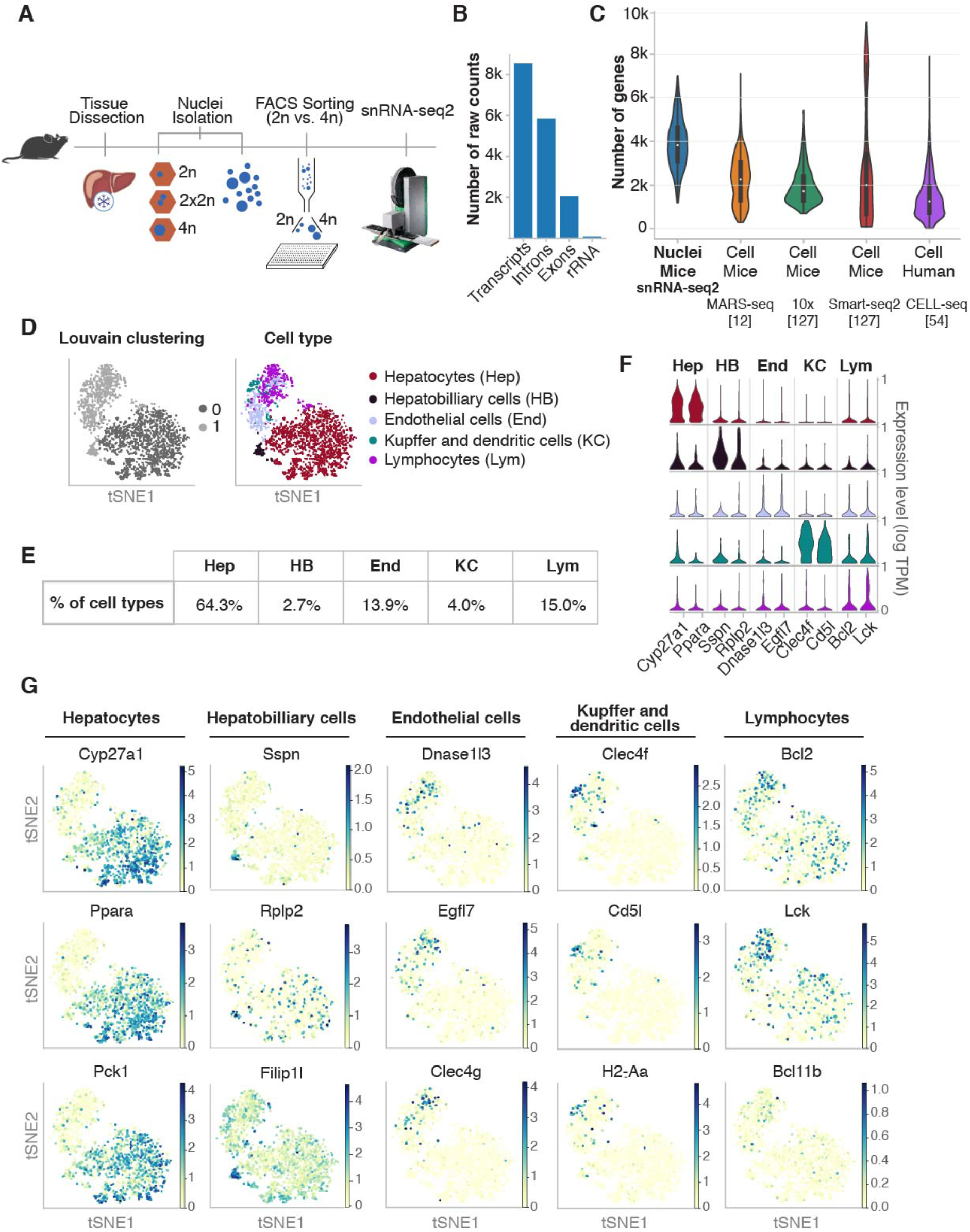
snRNAseq2 on frozen tissue identifies and characterize all major cell types present in liver. **A)** Outline of the pipeline used to perform snRNAseq2. Nuclei are isolated from flash frozen tissue, sorted by genome content in 384 well-plates and subjected to generation of full-length cDNA and sequencing. **B)** Quality control of sequenced data shows the number of raw counts per genome feature in the nuclear transcriptome. **C)** Violin plot comparing the high number of genes detected using snRNA-seq2 to publicly available single-cell RNA-seq datasets in which intact cells are isolated. **D)** t-SNE embedding of the nuclei by low resolution *Louvain* clusters (left), and annotated cell type (right) for all major cell types identified. **E)** Percentage of cell types identified by snRNA-seq2. **F)** Stacked violin plot depicting two illustrative marker genes per cell type. **G)** t-SNE colored by marker gene expression revealed cellular heterogeneity in all major cell population of the mouse liver.

The main improvement of our approach relies on the addition of a supplementary lysis buffer compatible with the generation of full-length cDNA and library preparation without the need for additional clean-up steps. This second lysis buffer (LB2) is compatible with a wide range of commercially available platforms and chemistries, including C1 Fluidigm (Fluidigm), well-plate approaches combined with liquid handling robots (Mosquito HV, TTP Labtech), as well as both SMARTer and NEBNext commercially available chemistries (Supp. Fig. S1D-E).

T-distributed stochastic neighbor embedding (t-SNE) was used to visualize nuclei FACS-sorted in an unbiased manner (Fig. 1D). Low resolution *Louvain* clustering showed that the nuclei were clustering into two groups (0, 1), corresponding to hepatocytes and non-hepatocyte cells (Fig. 1D, left). After higher resolution clustering and identification of the top differential expressed genes between clusters, we identified hepatocytes (64.3%), hepatobiliary cells (2.7%), kupffer and dendritic cells (4%), endothelial cells (13.9%) and lymphocytes (15.0%) as main cell clusters (Fig. 1D right and Fig. 1E; Supp. Fig. S1F-G; Supp. Table S1 and Supp. Table S2) [11, 12, 53, 54]. The expression distribution of key markers characteristic of those cell types showed that all liver cell populations can be identified with this methodology (Fig. 1F). Interestingly, visualization of key representative markers such as *Cyp2d26, Ppara, Pck1* for hepatocytes; *Sspn, Rplp2, Filip1l* for hepatobiliary cells; *Dnase1l3, Egfl7, Clec4g* for endothelial cells; *Clec4f Cd5l, H2-Aa* for Kupffer and dendritic cells; and *Bcl2, Lck, Bcl11b* for lymphocytes showed that transcriptional heterogeneity can be captured from frozen liver tissues for each cell type (Fig. 1G). Further in-depth characterization of liver cell populations revealed additional features related to their genome content and respective ploidy levels. Heat map of the top twelve differentially expressed genes showed that hepatocytes and hepatobiliary cells were mainly comprised by 2n and 4n nuclei while other cell types were primarily associated to 2n nuclei (Fig. 2A, and Supp. Fig. S2A).

**Figure 2.**
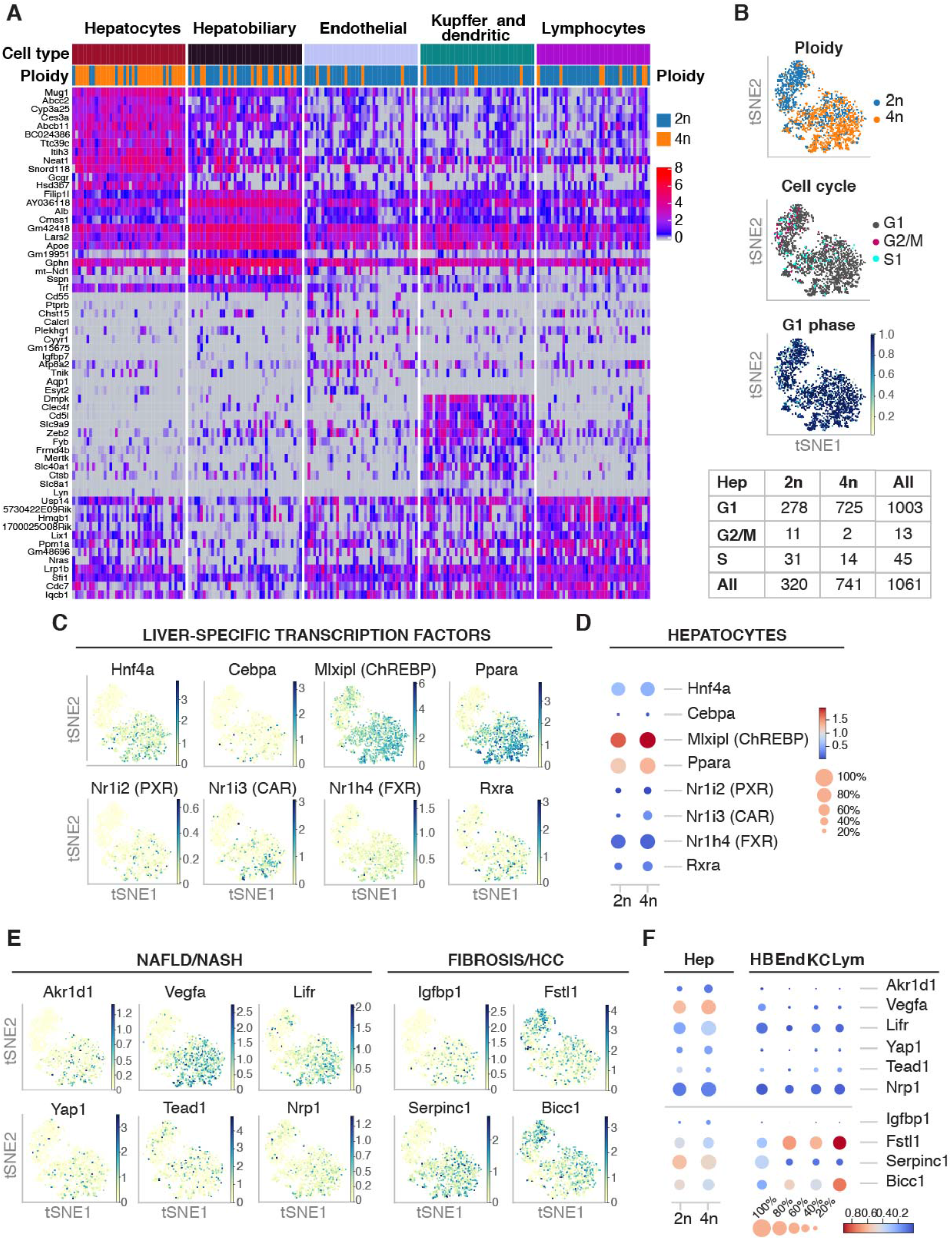
snRNA-seq allows deep profiling of single nuclei including key liver specific transcription factors and downstream target genes involved in healthy homeostasis and chronic liver disease. **A)** Heatmap showing the gene expression of the top twelve differentially expressed genes in forty randomly selected nuclei per cell type. Ploidy analysis shows that 4n nuclei are enriched in the hepatocyte cluster. **B)** Cell cycle analysis using *Cyclone* shows that the majority of nuclei are in G1 phase. t-SNEs colored by ploidy (top), assigned cell cycle phase (middle), and G1 score (bottom). Table showing the number of nuclei that are in each assigned phase for diploid and tetraploid nuclei from the hepatocyte cluster. **C)** t-SNE colored by the expression of key liver-specific transcription factors involved in liver homeostasis and hepatic function. **D)** Dot plot shows that the expression levels and the percentage of cells expressing key transcription factors can be dissected between 2n and 4n nuclei. **E)** t-SNEs colored by the expression of disease-related marker genes, separated into two main categories of liver disease (NAFLD/NASH and Fibrosis/HCC), showing cellular heterogeneity in different cell populations. **F)** Dot plot showing the expression of the disease-related marker genes across cell types in livers from young wildtype mice.

Consistent with the polyploid nature of hepatocytes [38, 55], we found as expected, that the hepatocyte cluster was enriched with a mixture of 2n and 4n nuclei (Fig. 2A). Nonetheless, few 4n nuclei were found in the non-hepatocyte clusters. Thus, we investigated whether cell cycle state could explain the presence of 4n nuclei in non-hepatocyte clusters (Fig. 2B and Supp. Fig. S2B-D) [44, 56, 57]. Cyclone was used to identify nuclei associated with G1, G2/M and S phase of cell cycle [57]. As expected in the liver tissue, the majority of nuclei were in G1 phase (1509 out of 1649) (Fig. 2B, Supp. Fig. S2B-D). This resulted in 94.5% of the hepatocyte cluster assigned to G1 phase, while 85.5% of nuclei in the non-hepatocyte cluster were in G1 phase. The number of 4n nuclei in the non-hepatocyte cluster was below 7%. This can be explained either by cell cycle stage (11 nuclei were in division) or technical bias during FACS sorting (e.g. nuclei tend to clamp together). Therefore, FACS sorting of nuclei according to their DNA content is a robust and accurate strategy to analyze hepatocytes with different levels of ploidy [38].

These results demonstrate that our method is highly sensitive to identify the main cell types in the liver and inspect the cellular heterogeneity present in intact frozen tissues.

### snRNA-seq2 uncovers transcription factors and rate limiting enzymes involved in chronic liver disease

In order to investigate whether the nuclear transcriptome can be used to address functional responses in the context of health and disease, we studied the expression levels of key liver-specific transcription factors (e.g. *Hnf4a, Ppara, Mlxpl*/ChREBP, *Cebpa*), nuclear receptors (e.g. *Rxa, Nrfl2/PXR, Nr1i3/CAR, Nrfh4/FXR*) and coactivators (e.g. *Ppargc1a/PGC1A,* Crebbp/CBP/p300, Ncoa1/SRC-1) [1–3, 6, 58] (Fig. 2C and Supp. Fig. S2E). Our approach allowed to deeply characterize the expression levels of upstream metabolic regulators in hepatocytes bearing two (2n) and four (4n) complete genomes. For the first time, we inspected the gene expression levels of key transcription factors and downstream effectors between 2n and 4n nuclei isolated from hepatocytes (Fig. 2D). For instance, Peroxisome proliferator-activated receptor α (*Ppara*) is the most abundant isoform expressed in hepatocytes with key roles in lipid metabolism and displays protective roles against Non-alcoholic fatty liver disease (NAFLD) [59–61]. *Ppara* expression was upregulated in 4n nuclei in comparison to 2n nuclei (Fig. 2D; Supp. Table S2 and Supp. Table S3). Similarly, Carbohydrate responsive element binding protein (ChREBP), encoded by the *Mlxpl* gene, is a carbohydrate-signaling transcription factor highly expressed in liver and adipose tissues and regulates the synthesis *de novo* of fatty acids [62–64]. *Mlxpl* is upregulated in 4n hepatocytes (Fig. 2D; Supp. Table S2 and Supp. Table S3). The precise interplay between upstream transcription factors and rate limiting enzymes in the hepatic lipid and glucose metabolism will determine the final outcome in chronic lipid overload and NAFLD. In consequence, investigating the functional differences between 2n and 4n hepatocytes is crucial to understand the development and progression of complex liver disease.

Recently, changes in the expression patterns of transcriptional regulators and downstream target genes have been associated with the development and progression of chronic liver diseases such as NAFLD [65, 66], non-alcoholic steatohepatitis (NASH) [67], fibrosis [15] and human hepatocellular carcinoma (HCC) [54]. Here we show that our methodology is highly sensitive to detect those gene markers in the nuclear transcriptome (Fig. 2E-F). In particular, dysregulation of *Akr1d1* has been associated with human NAFLD [66], and upregulation of *Wwtr1* (also known as TAZ) in human and murine NASH liver [68]. Furthermore, *Tead1* has been proposed as a marker of NASH in murine mouse models [69]. Additionally, *Vegfa*, *Lifr*, and *Nrp1* have been associated with the intrahepatic ligand-receptor signalling network involved in the pathogenesis of NASH [67] (Fig. 2E-F). Similarly, downregulation of *Bicc1* and *Fstl1* has been associated with advance fibrosis [70]. More recently, Aizarani *et al.* [54] have shown that the perturbation of gene signatures in individual cells was associated to HCC, for instance changes in *Serpin1c* and *Igfbp1* (Fig. 2E-F).

In summary, we are able to detect marker genes associated with chronic liver disease identified from bulk and single-cell transcriptomic analysis. This methodology has the potential to investigate previously archived frozen murine and human samples, and interrogate how changes in gene expression correlate with the development and progression of liver disease, taking into account cellular crosstalk and signaling pathways.

### snRNA-seq2 reveals differential transcriptional variability between 2n and 4n nuclei

Although recent single-cell transcriptomic studies have addressed cellular heterogeneity in the liver with respect to hepatocyte zonation [11, 12, 14, 15, 54], and the variability among non-parenchymal cells during chronic liver disease [67, 71–74], little is known about the cellular heterogeneity in hepatocytes with different levels of ploidy [55, 75–77].

Polyploidy occurs in the liver as normal tissue development [78], and is also associated with pathological conditions such as cancer and chronic liver disease [79, 80]. Importantly, ploidy increases with age [81–83], as well as in chronic liver conditions [84]. More specifically, the presence of tetraploid mononucleated hepatocytes has been associated with a poor prognosis in human hepatocellular carcinoma (HCC) [85, 86]. Accordingly, we further characterized the transcriptional profile of individual 2n and 4n nuclei from the hepatocyte cluster and investigated whether their transcriptomic profile differs among those two populations.

Across all cell types, the median number of genes detected in 4n nuclei was 1.36-times higher than in 2n nuclei (Supp. Fig. S3A and Methods). In particular, in the hepatocyte cluster a 1.25-times increase was detected in 4n compared to 2n nuclei (Fig. 3A). Therefore, genome duplication does not translate in doubling the number of detected genes, indicating that 2n and 4n nuclei might have different mechanisms of gene regulation to buffer gene dosage effects [87, 88] (Fig. 3A, Supp. Fig. S3A and Methods). Importantly, we saw that 2n and 4n nuclei from the hepatocyte cluster do not separate in t-SNE embedding based on their global gene expression profile (Fig. 3B); thus, they would be indistinguishable without a priori knowledge of their ploidy status. Herein, 2n and 4n nuclei from the hepatocyte cluster will be named 2n and 4n hepatocytes respectively.

**Figure 3.**
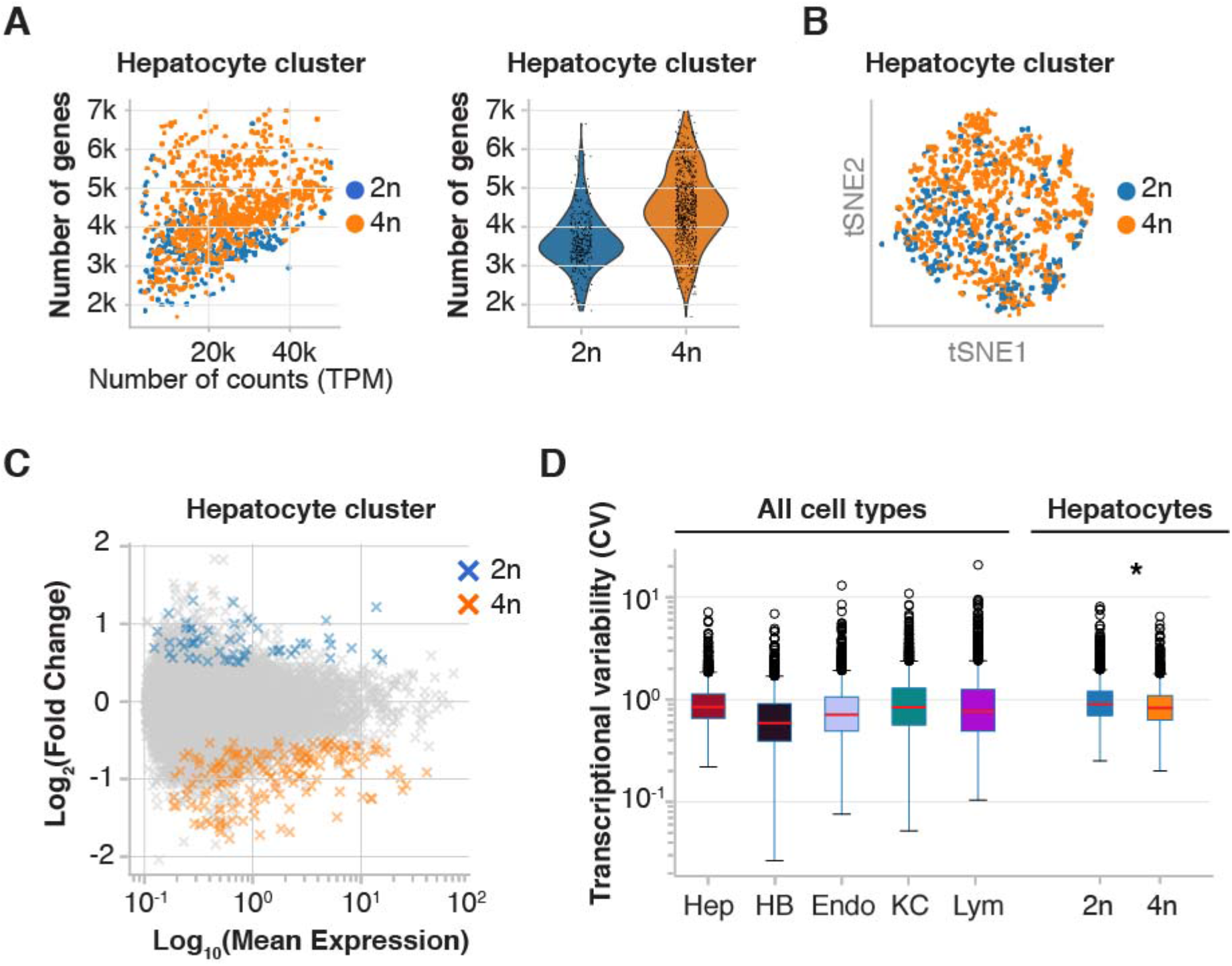
snRNA-seq2 reveals differences in gene expression between diploid and tetraploid hepatocytes. **A)** Scatterplot showing the higher number of genes per normalized counts detected in 4n hepatocytes compared with 2n hepatocytes (left), and violin plot showing that in 4n nuclei the number of genes detected is 1.25-time higher than in 2n hepatocytes (right). **B)** t-SNE embedding of the hepatocyte cluster shows that 2n and 4n nuclei cluster together. **C)** MA plot showing average logarithmically transformed mean expression (x-axis) *versus* the log2 fold change (y-axis) for pairwise comparison between 2n and 4n hepatocytes; 312 DEGs are depicted with crosses: 64 genes are upregulated in 2n nuclei (blue) and 248 genes are upregulated in 4n (orange). **D)** Box plot of the coefficient of variation shows that hepatobiliary cells present the lowest transcriptional variability across cell types (left), and that 4n hepatocytes are less variable than 2n hepatocytes (right).

Differential expression analysis showed that 2n and 4n hepatocytes are globally very similar. We detected 248 genes upregulated and 64 genes down regulated in 4n (Fig. 3C, Supp. Fig. S3B and Supp. Table S3). Subsequently, the transcriptional variability was estimated as coefficient of variation (CV) using log transformed data [89–91] (Methods). We confirmed that 4n hepatocytes showed a significant lower CV compared to 2n hepatocytes (2n are 1.09-times more variable than 4n, Mann–Whitney U test, p value 2.097e-14) (Fig. 3D). Interestingly, hepatobiliary cells showed the lowest CV compared with other cell types (Fig. 3D). Accordingly, the number of highly variable genes (HVGs) was higher in the 2n hepatocytes compared to the 4n hepatocytes (Supp. Fig. S3C and Supp. Table 4).

In summary, 2n and 4n hepatocytes showed a similar transcriptional profile in young mice, however 312 DEGs were detected between these two cellular states. Importantly, the number of cells contributing to the DEG were different between 2n and 4n hepatocytes. These results revealed the concealed cellular heterogeneity present in hepatocytes with different levels of ploidy.

### Tetraploid nuclei show extensive co-expression of liver stem cell markers

Polyploid hepatocytes have been associated with terminal differentiation and senescence [40]. However, a growing body of evidence indicates that polyploid hepatocytes retain their proliferative potential [77, 92–95]. To further analyze the functional characteristics of 2n and 4n hepatocytes, Gene Ontology (GO) analysis was first performed on significant DEGs (Supp. Fig. S4A, Supp. Table 3). DEG in 2n hepatocytes were only enriched in one category (i.e. Small molecule metabolic process), while DEG in 4n hepatocytes were enriched in twenty categories (Supp. Fig S4A). Secondly, we extended our GO analysis to the top hundred most differentially expressed genes (Fig. 4A and Supp. Table 3). Interestingly, comparisons between 2n and 4n hepatocytes showed that 4n hepatocytes were further enriched in pathways involved in lipid, cholesterol and xenobiotic metabolism (Fig. 4A and Supp. Table 3).

**Figure 4.**
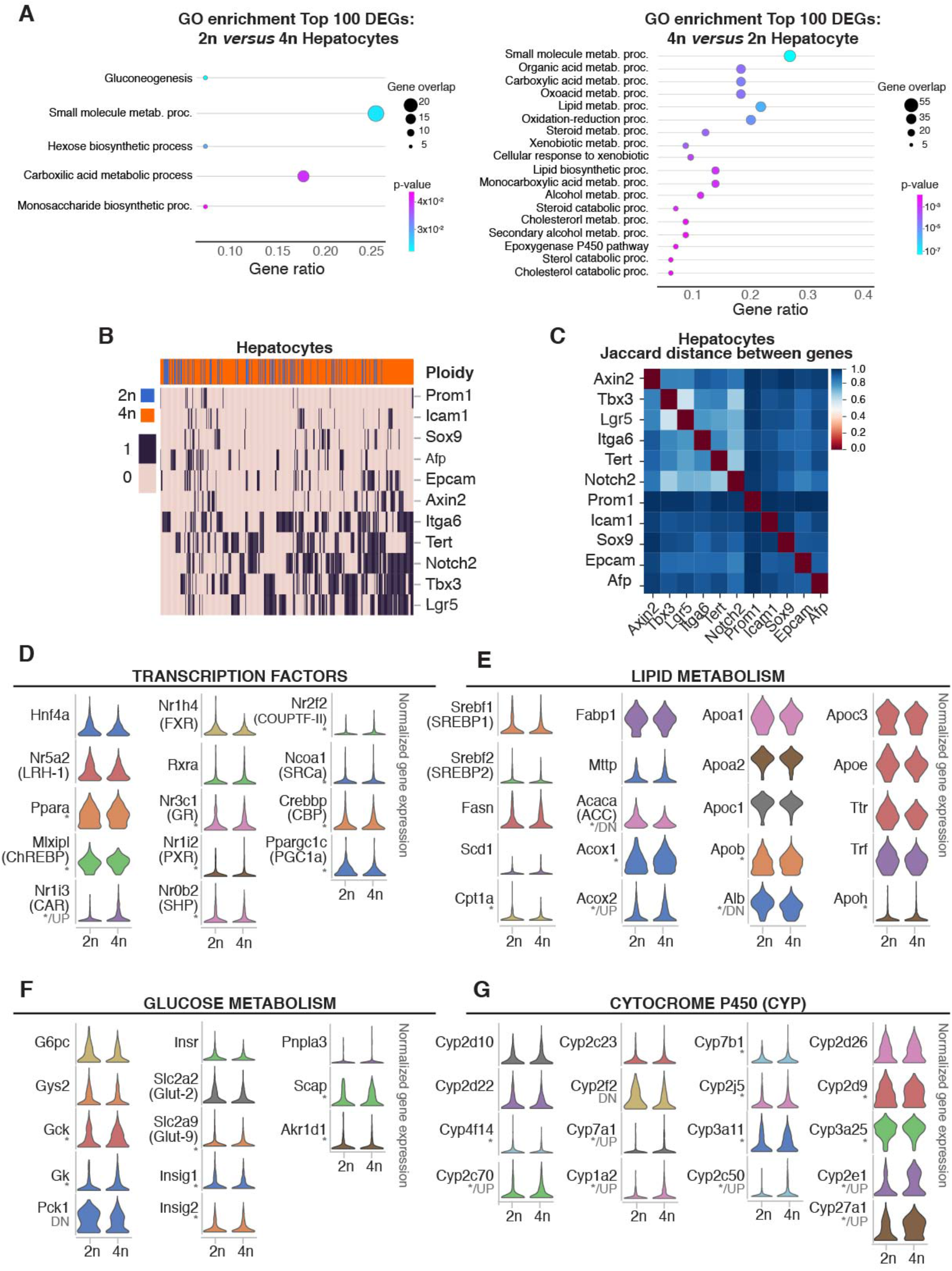
Functional characterization of 2n and 4n hepatocytes at the single-cell level. **A)** Gene set enrichment analysis of genes upregulated in 2n hepatocytes in comparison to 4n hepatocytes (left), and *vice versa* (right). **B)** Heatmap showing extensive co-expression of stem cell markers in hepatocytes (binary expression: 1-detected, 0-non-detected). Ordering of the cells and genes is based on hierarchical clustering between nuclei (columns), and stem cell markers (rows), revealing the stem cell markers are co-expressed in 2n and 4n hepatocytes. **C)** Heatmap showing pairwise Jaccard distances between stem cell markers, reveals one main module of co-expression in hepatocytes. **D)** Violin plots showing changes in gene expression distribution in key liver-specific transcription factors; **E)** Violin plots showing key regulator of the hepatic lipid metabolism; **F)** Violin plots showing key regulators of hepatic glucose metabolism, and **G)** Violin plots of representative genes from the cytochrome P450 family involved in the metabolism of xenobiotics in 2n and 4n hepatocytes (*: significant changes in gene expression distribution, UP: significant upregulation in 4n, DN; significant downregulation in 4n).

In order to further characterize the basal differences between 2n and 4n hepatocytes, we studied the regenerative and proliferative properties of these two cellular states. It has been suggested that diploid hepatocytes, located in proximity to the central vein of the liver lobule, act as stem cells during homeostasis and in response to injury [96]. However, it has recently been shown by multicolor reporter allele system that polyploid hepatocytes proliferate in chronically injured livers [77, 92, 97]. Independent studies by Su *et al.* using AXIN2 lineage tracing have shown that hepatocytes upregulate AXIN2 and LGR5 after injury throughout the liver lobe [77, 92, 97]. To investigate if there are specific subpopulations of cells with stem cell properties enriched in 2n or 4n hepatocytes, several stem/progenitor cell-like marker genes were selected, including *Icam1, Afp, Sox9, Epcam, Axin2, Tbx3, Itga6, Tert, Lgr5* and *Notch2* [76] (Fig. 4B and Supp. Fig. S4B). Five of these markers, *Notch2, Tbx3, Itga6, Lgr5* and *Tert* were expressed in more than 40% of the cells analyzed (Supp. Fig. S4C). We observed that both 2n and 4n hepatocytes expressed several stem/progenitor markers and we did not find an enrichment of those genes in diploid hepatocytes (Fig. 4B and Supp. Fig. S4C). In order to investigate whether these markers were co-expressed in the same nuclei, the Jaccard index was used to measure the probability of those genes being co-expressed in the same cell/nucleus [98, 99]. In particular, the Jaccard distance measures dissimilarity between genes, and lower distance indicate higher probability of co-expression (Fig. 4C). With the Jaccard distance one module was identified for *Afp*, *Epcam*, *Sox9*, and *Icam1* (Fig. 4C and Methods). This module showed pair-wise similarity independently of gene expression levels. Additional genes showed higher distance and lower probability of co-expression*: Axin2*, *Tbx3*, *Lgr5*, *Itga6*, *Tert*, and *Notch2* (Fig. 4C and Methods). Furthermore, the percentage of nuclei expressing those gene markers was similar in both 2n and 4n hepatocytes (Supp. Fig. S4C) indicating that 2n and 4n hepatocytes have similar stem cell properties. Interestingly, more than two stem/progenitor markers were co-expressed in both 2n and 4n hepatocytes (Fig. 4B, and Supp. Fig. S4D; Methods) strongly supporting the notion that polyploid hepatocytes have regenerative potential.

These results show that in young livers, there is no subpopulation of diploid hepatocytes enriched in stem/progenitor markers, and that polyploid hepatocytes co-express genes associated to proliferative and regenerative functions, and therefore have the potential to contribute to organ regeneration and liver homeostasis.

### Changes in expression pattern distribution in 4n nuclei indicates a high capacity for adaptation and regeneration

A focused analysis on energy homeostasis also revealed changes on the distribution pattern of key metabolic genes involved in lipid and glucose metabolism (Fig. 4D-G, Supp. Fig. S5 and Supp. Table 5). Some liver-specific transcription factor showed no change in mean expression or distribution pattern between 2n and 4n hepatocytes, for instance, *Hnf4a, Nr5a2* (LRH-1), *Nr1h4* (FXR), and *Rxa* (Fig. 4D). However, *Mlxipl* (ChREBP), *Nr3c1* (GR), *Nr1i2* (PXR), *Nr0b2* (SHP), *Nr2f2* (COUPTF-II), and the coactivators *Ncoa1* (SRCa), *Crebbp* (CBP), and *Ppargc1c* (PGC1a) showed a different distribution pattern between 2n and 4n but no changes in mean expression. In some cases, such as *Nr1i3* (CAR), changes in distribution were associated with upregulation of its expression levels in 4n hepatocytes (Fig. 4D and Supp. Table 3 and Supp. Table 5). We further studied critical regulators of lipid metabolism and found that *Cpt1, Acox1, Apob* and *Apoh* showed changes in distribution pattern but no changes in mean expression. Albeit, in *Acaca* (ACC) and *Alb*, we found that their distribution pattern changed and its expression was downregulated in 4n hepatocytes, while *Acox2* was upregulated in 4n.

Regarding the glucose metabolism, changes in the distribution pattern of *Gck, Gk, Slc2a9* (Glut-9), *Insig1, Insig2, Scap,* and *Akr1d1* were found (Fig. 4F, Supp. Fig. S5 and Table 5). Interestingly, we observed considerable changes in the metabolism of xenobiotics and cytochrome P450 superfamily (Fig. 4G, Supp. Fig. S5 and Supp. Table 5). For instance, *Cyp4f14, Cyp2j5, Cyp2d9,* and *Cyp3a25* showed changes in gene expression distribution, while *Cyp2c70, Cyp7a1, Cyp1a2, Cyp2c50, Cyp2e1,* and *Cyp27a1* were additionally upregulated in 4n hepatocytes (Fig. 4G).

In summary, rate limiting enzymes involved in energy homeostasis and metabolism of drugs showed a significant change in expression distribution between 2n and 4n hepatocytes. Changes in gene expression distribution have been associated to genetic plasticity and higher adaptation [100–103], which could be determinant when the liver is faced with a chronic or overwhelming insult.

### Hepatic metabolic zonation determines gene expression levels independently of the ploidy status

Hepatocytes and endothelial cells have been spatially located in the liver along the portal-central axis of the liver lobule [11, 12]. Recently, 2n and 4n hepatocytes have been localized to specific metabolic zones (periportal or pericentral) or randomly interspersed throughout the hepatic lobe [96, 104, 105]. We used diffusion pseudotime (dpt), visualized in diffusion maps, to infer the pseudospatial ordering of nuclei according to defined markers associated with liver zonation and their metabolic specialization [12, 106, 107] (Fig. 5A and Supp. Fig. S6A). *Louvain* clustering on these zonation markers established five clusters (Supp. Fig. S6A). After visual investigation of pericentral and periportal marker genes on the diffusion map (Fig. 5C and Supp. Fig. S6B), three clusters (0, 2, and 3) were aggregated into a periportal cluster, and two clusters (1 and 4) into a pericentral cluster (Fig. 5C and Supp. Fig. S6B). Then, the percentage of 2n and 4n hepatocytes that were in the pericentral (CV) and periportal (PV) clusters were calculated respectively (Fig. 5B and Methods) [12]. We observed a 1.3-times relative enrichment of 4n hepatocytes in the pericentral cluster, indicating that they reside close to the central vein. This is in agreement with recent studies in which polyploid hepatocytes have been preferably located by histological analysis in the pericentral zone, and diploid hepatocytes in the periportal zone using cell-lineage tracing [77, 108].

**Figure 5.**
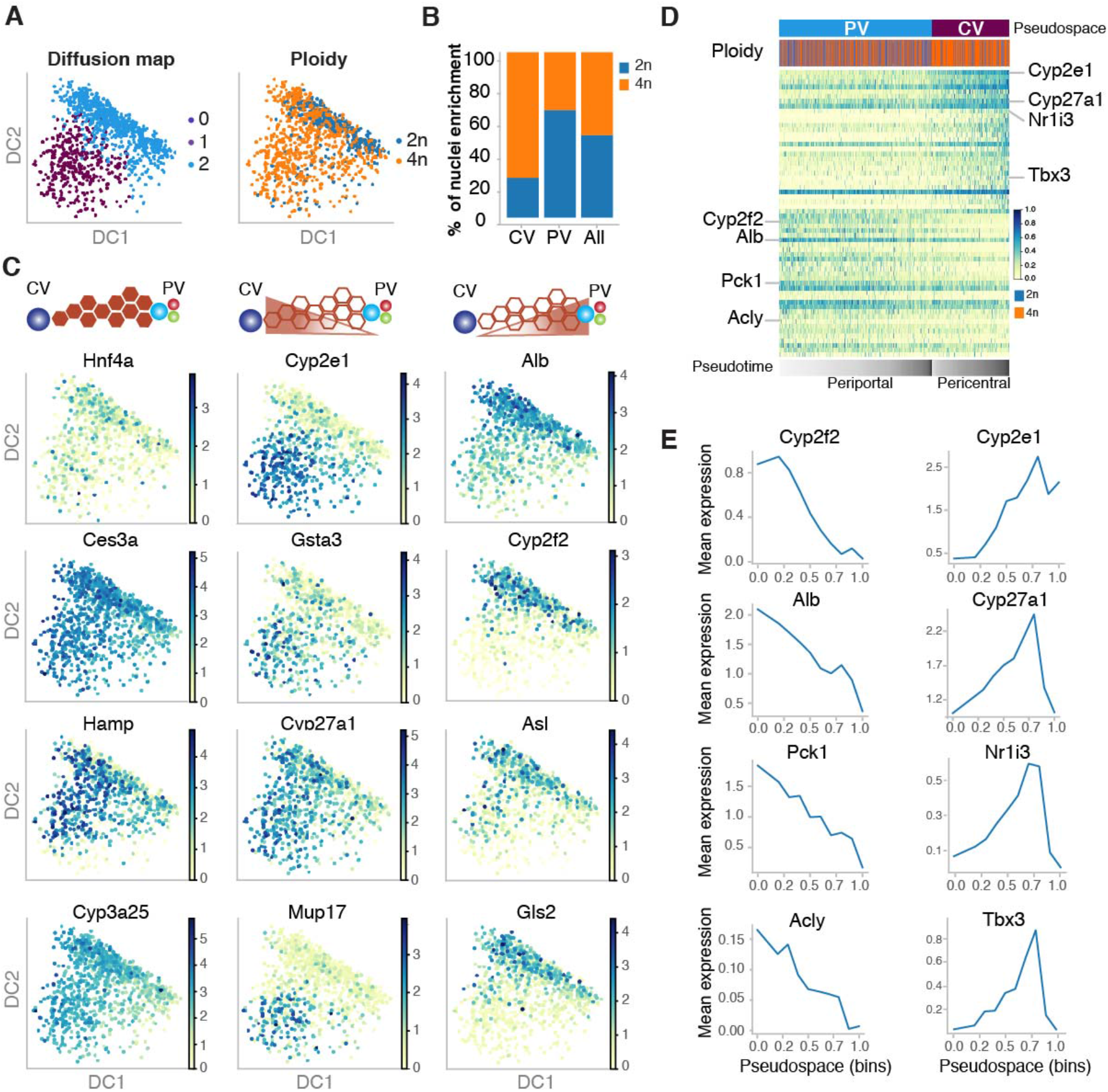
Pseudospatial ordering of hepatocytes along the liver lobe shows functional crosstalk between ploidy and zonation. **A)** Diffusion map based on zonation markers, colored by *Louvain* clusters (left) and ploidy (right). **B)** Bar plot showing higher percentage of nuclei in the pericentral cluster in comparison to the periportal cluster, colored by ploidy. **C)** Diffusion maps colored by the expression of zonation marker genes, for representative non-zonated genes (*Hnf4a, Ces3a, Hamp, Cyp3a25*), pericentral genes (*Cyp2e1, Gsta3, Cyp27a1, Mup17*) and periportal genes (*Alb, Cyp2f2, Asl, Gls2*). **D)** PAGA path heatmap showing the top 30 differentially expressed genes per cluster between the pericentral and the periportal cluster of nuclei, showing an enrichment of 4n hepatocytes in the pericentral zone. **E)** Line plots depicting mean expression of individual zonation markers per bin of the diffusion pseudospace vector for representative genes enriched in the pericentral or periportal zone respectively.

We further studied the expression levels of classic markers of liver zonation such as *Cyp2e1, Gsta3, Cyp27a1* and *Mup17* for the central vein, and *Alb*, *Cyp2f2, Asl,* and *Gls2* for the portal vein (Fig. 5C, Supp. Fig. S6 and Supp. Fig. S7). Non-zonated markers were also selected, for instance *Hnf4a* which mRNA expression is non-zonated as opposed to its protein levels [10]. Additional non-zonated genes including *Ces3a, Hamp,* and *Cyp3a25* were also studied (Fig. 5C, Supp. Fig. S6 and Supp. Fig. S7). In order to verify the spatial distribution of 4n hepatocytes, we binned the vector of diffusion pseudotime into 10 bins and visualized the mean expression of specific zonation markers along these bins (Fig. 5C and Supp. Fig. S7). This approach is very powerful to visualize simultaneously the decrease and increase in mean expression of periportal and pericentral genes along these bins. These results showed that liver zonation can be investigated by diffusion pseudotime using our snRNA-seq2 approach (Fig. 5C, Supp. Fig. S6 and Supp. Fig. S7, Table S6). Moreover, we generated a partition-based graph abstraction (PAGA) to visualize coordinated changes in expression across the pseudospace in the hepatic lobe [109] (Methods). The thirty most differentially expressed genes in each of the two zonation clusters were selected along the diffusion pseudotime, again showing an enrichment of 4n hepatocytes in the pericentral cluster and the expected changes of pericentral and periportal markers, respectively (Fig. 5D-E, Supp. Fig. S6 and Supp. Fig. S7). Out of 224 zonation markers that were upregulated in the pericentral cluster, 55 were upregulated only in 4n hepatocytes. Meanwhile, out of 68 zonation markers that were upregulated in the periportal cluster only 12 were upregulated in 2n hepatocytes.

In summary, our results indicate a tight coordination between gene dosage and spatial location within the liver lobe. Therefore, hepatocytes adjust their gene expression according to their spatial distribution showing a functional crosstalk between liver zonation and ploidy.

## Discussion

Single-cell genomics allows the unbiased exploration of cell states and cell types at single-cell resolution, leading to a revolutionary change in our understanding of liver biology and disease pathogenesis [16]. However, single-cell RNA-seq (scRNA-seq) in the liver is associated with some caveats. First, dissociation protocols including enzymatic or mechanical dissociation lead to changes in the cellular transcriptome [17–19]. Secondly, dissociation may lead to underrepresentation of certain cell types due to cell fragility or large cell-size, such as hepatocytes [110]. Thirdly, it relies on the isolation of intact cells from fresh tissues, which hinders its clinical application on human samples. In this context, single-nucleus RNA-seq has emerged as a complementary approach that relies on the unbiased assessment of nuclei from all cells present in a tissue [20, 21, 36]. The analysis of the nuclear transcriptome has been proven to be very powerful to study cell type diversity in the mouse and human tissues, including brain [21, 23–27, 111]; spinal cord [50]; breast cancer [51]; kidney [29–32]; lung [28]; heart [33, 34]; and a variety of human tumor samples [36].

We have developed a single-nucleus RNA-seq2 method that improves the lysis of the nuclear membrane and significantly increases the number of transcripts detected per nuclei, even when considering an intact cell (Fig. 1 and Supp. Fig. S1). Our method is highly sensitive, reproducible and it has been specifically designed for archived liver samples (Fig. 1). The unbiased sorting of single-nuclei allows to study all the main cell types comprised in the liver, and it can be applied to interrogate cellular crosstalk and transcription factor networks (Fig. 1 and Fig. 2). Indeed, snRNA-seq2 has the potential for comprehensive analysis of fresh and archived flash frozen liver samples from both mice and humans.

Notably, scRNA-seq has revealed the high degree of functional specialization of cell types within the liver lobe, for instance for hepatocytes, endothelial cells [9, 11, 12, 54], and macrophages [14, 15, 71]. Additionally, it has been shown how changes in mRNA highly correlate with changes at the protein levels in the liver [10]. However, little is known about the functional role of polyploid hepatocytes at the single-cell level. Recently, Katsuda *et al.* have shown in rats how genes associated to hepatic zones have differential expression pattern in polyploid hepatocytes [76]. Here, we present a genome-wide analysis of the nuclear transcriptome of individual 2n and 4n nuclei in mice (Fig. 3-5). Our study is focused on the analysis of ploidy based on the DNA content of each nucleus (2n and 4n), and we cannot resolve whether 2n nuclei belong to either diploid cells or binucleated tetraploid cells (2nx2) [40, 80]. Moreover, it has been shown that the hepatocyte volume does not depend on the number of nuclei but rather on their polyploidy status, whereas mono-nucleated hepatocytes (2n) and bi-nucleated hepatocytes (2nx2) have the same cellular volume [8, 112]. In young wildtype mice, an incomplete cytokinesis leads to tetraploid cells (2nx2) that is generally associated to fidelity of chromosome transmission [40, 80]. For these reasons, we have focused our analysis on the 2n and 4n nuclei, for which we do not expect chromosomal abnormalities [113, 114].

Our transcriptomic analyses revealed that 2n and 4n hepatocytes are globally very similar, but 312 genes are found to be differentially expressed between them (Fig. 3). This suggests that if polyploid hepatocytes increase during pathological processes, they could lead to gene expression imbalances of functionally relevant genes (Fig. 3 and Fig. 4). Additionally, the transcriptional variability in young mice is also lower in 4n hepatocytes, which it has been shown previously by means of single molecule fluorescence in situ hybridization (smFISH) for the beta-actin (*Actb*) gene [88]. Whether transcriptional variability changes during ageing [46] or in chronic liver disease still remains to be investigated. Interestingly, the presence of mononucleated tetraploid hepatocytes has been associated with human hepatocellular carcinoma (HCC) [39, 84, 85, 115]. Likewise, the number of polyploid hepatocytes also increases in models of NAFLD, including ob/ob mice and wild-type mice fed with methionine-choline-deficient diet (MCD) or high-fat diet (HFD) [84, 116]. Therefore, the altered number of 2n and 4n hepatocytes is strongly associated with NAFLD in both animal models and patients. The transcriptomic analysis of 2n and 4n hepatocytes in bio archived samples could open an entirely new strategy to understand disease pathogenesis and search for new therapies to treat liver disease.

The liver is characterized by its regenerative potential. However, the cellular origin that triggers liver homeostasis and repair is still under debate. Recently, complementary studies using different models of cell lineage-tracing have shown that all hepatocytes have comparable self-renewal potential [77, 92, 97]. Our transcriptomic analysis also supports the notion that the vast majority of hepatocytes, regardless their ploidy level and location within the hepatic lobe, co-expresses stem/progenitor gene markers and have the potential to contribute to liver homeostasis after hepatic injury (Fig. 4B-C).

Furthermore, the broad spatial heterogeneity in the liver shows that key liver functions are zonated [9], and recently it has been shown that the zonation can be perturbed during liver fibrosis [14, 15]. In this context, the spatial distribution of polyploid hepatocytes and its functional consequences are still in debate [8, 40, 77, 85, 108]. We used diffusion pseudotime to infer the pseudospatial ordering of 2n and 4n hepatocytes according to defined markers of liver zonation, and we found that 4n nuclei are distributed throughout the hepatic lobe with an enrichment in the pericentral zone (Fig. 5). These results support previous studies in which polyploid hepatocytes are quantified by lineage-tracing [77] or smFISH for the glutamine synthetase (*Glu1*) [108]. In summary, we have found that 4n hepatocytes are enriched 1.3-times in the pericentral zone, and that ploidy and liver zonation are tightly regulated. Altogether, we have discovered that the division of labor in hepatocytes is linked to both their spatial distribution and ploidy levels, and this crosstalk could be affected during ageing and chronic liver disease.

## Conclusion

In summary, our methodology has the potential to investigate previously archived frozen murine and human samples, and interrogate how changes in gene expression correlates with the development and progression of liver disease, taking into account cellular crosstalk and signaling pathways. Our analysis shows that hepatocytes in young livers, without injury or selective pressure, express stem cell markers across the portal-central axis independently of their ploidy status. These findings strongly support recent reports showing that polyploid hepatocytes are proliferative and contributors to liver regeneration. Importantly, hepatocytes with different levels of ploidy adjust their gene dosage according to their position in the liver, showing a functional crosstalk between liver zonation and ploidy. We anticipate that changes in the number of 2n and 4n hepatocytes in the overall liver cell composition will lead either to protective or deleterious outcomes during ageing and chronic liver diseases.

## Methods – Experimental methods

### Mice and Liver tissue collection

All wild-type C57BL/6 mice were purchased from Charles River UK Ltd (Margate, United Kingdom) and were maintained under specific pathogen-free conditions at the University of Cambridge, CRUK – Cambridge Institute under the auspices of a UK Home Office license. These animal facilities are approved by and registered with the UK Home Office. All male C57BL/6 animals were sacrificed by an approved scientist in accordance with Schedule 1 of the Animals (Scientific Procedures) Act 1986. Young 3 month old male mice were sacrificed and individual pieces of the harvested liver were immediately flash-frozen. A small piece of liver tissue was collected in 4% paraformaldehyde. All young animals for this study were macroscopically inspected for any signs of pathologies or abnormalities.

### Single Nuclei Isolation / Homogenization

For the single nuclei isolation, the protocol described by Krishnaswami *et al*. [21] was followed with several modifications. In summary, fresh frozen liver tissues (~3 mm^3^) were homogenized using a 2 mL Dounce homogenizer (Lab Logistics, #9651632) in 1 mL of ice cold Homogenization Buffer, HB (250 mM sucrose, 25 mM KCl, 5 mM MgCl_2_, 10 mM Tris buffer, 1 μM DTT, 1 x protease inhibitor tablet -Sigma Aldrich Chemie), 0.4 U/μL RNaseIn (Thermo Fisher Scientific), 0.2 U/μL Superasin (Thermo Fisher Scientific), 0.1% Triton X-100 (v/v) and 10 μg/mL Hoechst 33342 (Thermo Fisher Scientific) in RNase-free water.

Before the dounce homogenization, the frozen tissue was placed in a pre-chilled petri dish on ice containing 1mL of HB and finely cut with a pre-chilled scalpel blade until all the pieces can be transferred to the homogenizer using a wide orifice 1 mL tip. Throughout the procedure all pipetting steps of transferring the samples were made using wide orifice tips (Rainin, #17014297) to minimize shear force.

Always on ice, we performed 5 very slow strokes with the loose inner tolerance pestle and 10 more strokes using the tight inner tolerance pestle. Collecting 10 μL of the solution, the nuclei suspension was inspected under a microscope on a Neubauer chamber with 10 μL of Trypan blue (0,4%). If aggregates were predominant, up to 5 more strokes were performed using the tight pestle. The suspension was passed through a 50μm sterilized filter (CellTrics, Symtex, #04-004-2327) into a 5mL Eppendorf pre-chilled tube washing the Dounce homogenizer with additional 500 μL of cold homogenization buffer. Subsequently, the suspension was centrifuged at 1,000g for 8 min in an Eppendorf centrifuge at 4°C (5430R). The pellet was then resuspended in 250μL of pre-chilled homogenization buffer. To ensure high quality of single-nuclei suspensions, we performed a density gradient centrifugation clean-up for 20 minutes, using Iodixanol gradient (Optiprep, D1556, Sigma Aldrich Chemie). The final pellet was resuspended gently in 200μL of nuclei storage buffer (NSB) (166.5 mM sucrose, 5mM MgCl_2_, 10mM Tris buffer pH 8.0) containing additional RNase inhibitors 0.2 U/μL Superasin (Thermo Fisher Scientific, #AM2696) and 0.4 U/μL Recombinant RNase Inhibitor (Takara Clontech #2313A). A final visual inspection and counting was performed under a microscope and the single nuclei suspension was filtered through a 35μm cell strainer cap into a pre-chilled FACS tube prior FACS sorting.

### Flow cytometry

Hoechst dye was used to stain all nuclei during nuclei isolation, allowing to distinguish between diploid and tetraploid nuclei using a FACS sorter (BD FACSAria Fusion 1 and/or BD FACSAria 3) with a 100μm nozzle. Before sorting, 384-well thin walled PCR plates (BioRad, #HSP3901) were freshly prepared with 940nL of Reaction Buffer (following manufacturer instructions, only 1uL of 10X Reaction Buffer is diluted in 2,75μl of water). Herein, this reaction buffer will be termed Lysis Buffer 1-LB1) (Takara kit SMART-Seq v4 Ultra Low input RNA) using a liquid miniaturization robot (Mosquito HV, STP Labtech) and kept on ice. Reaction buffer was prepared following the manufacturer instruction adding 1μL of RNAse Inhibitor in 19μL of 10X Lysis Buffer. The FACS droplet delay and cut-off point was optimized prior to every sorting, the plate holder was cooled at 4°C and all settings and calibrations were done by the FACS operator while samples were processed to avoid additional delays that could lead to RNA degradation of the samples. The gating strategies used were described elsewhere [38]. The sorting accuracy in the 384 well-plates was assessed using a colorimetric method with tetramethyl benzidine substrate (TMB, BioLegend, #421501) and 50 μg/ml of Horseradish Peroxidase (HRP, Life Technologies, #31490) [117]. Only when the plate alignment testing results to single events to more than 95% of the wells, we proceed with the fresh isolated nuclei. The nuclei suspended in NSB was diluted at ~10x 10^4^ nuclei per mL and kept on average 200 to 1000 events per second as flow rate. In the sorting layout, we sorted diploid (2n) cells in half of the 384 well plate and tetraploid (4n) in the remaining half for each individual. After sorting, every plate was sealed (MicroAmp Thermo Seal lid, #AB0558), shortly vortexed (10 seconds), centrifuged (prechilled at 4°C, 2000g for 1 minute), frozen on dry ice and stored at −80°C, until cDNA synthesis.

### snRNAseq-2

For the generation of double stranded full-length cDNA we used the Smart-Seq2 chemistry from a commercial kit (SMART-Seq v4 Ultra Low input RNA, Takara) and we optimized the lysis of the nuclear membrane by adding a supplementary lysis buffer (Lysis Buffer 2, LB2). Firstly, using a low volume liquid handling robot, we miniaturized the volumes reducing the amount 4-times and keeping the same ratios for all the reagents, significantly lowering the experimental cost. Liquid handling robots allowed to increase pipetting accuracy, maintained a sterile and temperature controlled environment and reduced the user variability and potential cross-contamination. Our additional lysis buffer 2 (LB2) consist in a mixture of 0.4 % NP-40 (v/v) final concentration (Life Tech, #85124) and 0.1% Triton-X100 (v/v) final concentration (Fisher, #10671652). Using the Mosquito HV (Labtech STP), we added 2190nL of our additional snRNAseq-2 lysis mix in each well together with the first strand synthesis primers and the spike-in controls (ERCC). Ratios to the final volume (3.125μl) of the total Lysis Buffer (1 and 2) are shown in parentheses (NP40 2% (2.5 / 12.5, Triton-X100 1% (1.25 / 12.5), ERCC spike-in at 1/300.000 dilution (1 / 12.5) and 3’ SMART-seq CDS Primer II A (2 / 12.5) and additional water (2 / 12.5). Every plate was thawed directly on a −20°C chilled holder while LB2 was added by the Mosquito HV robot. Then, the plate was sealed, vortexed vigorously (20 seconds in a Mixmate (Eppendorf)–2000 rpm), centrifuged (30 seconds in a pre-chilled Eppendorf 5430R at 4oC, 2000g) and placed in a Thermal cycler (BioRad C1000) for 6 minutes at 72°C). We suggest to use the same lot (#00769049) of ERCCs (Life Technologies, #4456740) per project. ERCC spike-ins were dilute 1 in 10, in water with 0.4 U/μL Recombinant RNase Inhibitor (Takara Clontech #2313A), aliquoted, and stored at −80oC. Fresh dilution of 1 in 300,000 and 1 in 100,000 were prepared immediately before the first strand synthesis.

Next, reverse transcription and Pre-PCR amplification steps were followed as described by the manufacturer. We maintained our 4-times reduced volumes for all steps. As a final modification on the Takara kit protocol, we optimized the PCR cycles program for the cDNA amplification. We used 21 cycles and a PCR programs consisting on: 1 min at 95°C, [20 sec at 95 °C, 4 min at 58 °C, 6 min at 68 °C] x 5, [20 sec at 95 °C, 30 sec at 64 °C, 6 min at 68 °C] x 9, [30 sec at 95 °C, 30 sec at 64 °C, 7 min at 68 °C] x 7, 10 min at 72 °C).

Internal ERCC spike-ins were used as positive controls and the cDNA yield was assessed in an Agilent Bioanalyzer with a High Sensitivity DNA kit. In our protocol we did not performed a bead clean-up before final library preparation. The miniaturization details and Mosquito programs used can be found in the supplementary information.

### RNA-seq library preparation and sequencing

Sequencing libraries were prepared using the standard Illumina Nextera XT. DNA Sample Preparation kit (Illumina, #FC-131-1096) and the combination of 384 Combinatorial Dual Indexes (Illumina-Set A to D, #FC-131-2001 to FC-131-2004). Using the Mosquito liquid handling robot, the Nextera XT chemistry was miniaturized reducing 10-times the manufacturers volume as previously described [118, 119] (Supp. Table 7). For the library preparation 500nL of the undiluted cDNA was transferred in a new 384 well-plate containing 1500nL of Tagmentation Mix (TD and ATM reagents). Accordingly, all Nextera XT reagents (NT, NPM and i5/i7 indexes) were added stepwise to a final library volume of 5μL per well. We prepared a 384 well-plate with the combination of the i5/i7 index adapters, which was used in our miniaturization protocols. The final PCR amplification was 12 cycles. After library preparation, 500nL from each well was pooled together to a final volume of ~192μl to perform a final AMPure XP bead (Beckman Coulter, #A63882) clean-up. For our single nuclei libraries, two consecutive clean-ups with a ratio of sample to bead 0,9X led to library sizes between 200 and 1000bp, with no trace of adaptors. The final libraries were assessed using a HS DNA kit in the Agilent Bioanalyzer.

### Sequencing

All the final 384-pooled libraries were sequenced using Illumina HiSeq4000 NGS sequencer in a paired-end 150 bases length. Each 384-pooled library was sequenced in one lane leading on average to ~1 million reads per nuclei.

### Liquid handling robots and miniaturization

As described before in the snRNAseq2 section, the use of automated liquid handler as the Mosquito HV, allowed the fast and accurate processing of 384 well-plates. For each of the steps in the cDNA synthesis and the library preparation a ‘source’ plate (384-well Low Volume Serial Dilution -LVSD, STP Labtech) was used for reagents. Importantly, the over-aspirating option must be switched off to minimize the ‘dead’ reagent volume. Mosquito HV was placed in a flow cabinet maintained in pre-PCR and RNase and DNA free room. The detailed protocols for the Mosquito HV can be found in supplementary information.

## Methods - Computational methods

### Read alignment and pre-processing

Raw sequencing reads were mapped against a customized genome containing both mm10 (GRCm38, assembly version 93), and the ERCC92 sequences [120] . Mapping was done using STAR-2.7.1a with the following parameters --*outFilterMultimapNmax 1 --outSAMtype BAM SortedByCoordinate*. Potential PCR-duplicates were identified and removed using*MarkDuplicates* from*picard tools version 2.20.2* with the option *REMOVE_DUPLICATES=true.* For every single nucleus, reads mapping to individual transcripts were counted and summed per gene using htseq-count version 0.11.3 with the following parameters: *-m intersection-nonempty -f bam -r pos -s no --nonunique all -t transcript -i gene_id --additional-attr=gene_name.*

The resulting raw count matrix was loaded into python and stored as an *AnnData* object, *anndata version 0.7.1.* Downstream analysis pipeline was inspired by following the tutorial from Luecken *et al.* [5]. Unless described otherwise, functions implemented in *scanpy* (version 1.4.5.2.dev6+gfa408dc7) were used in downstream analysis [109]. Additionally to the count matrix based on full transcripts, count matrices for exonic and intronic reads, respectively were built using featureCounts [121].

### Quality control and filtering

Starting with a matrix of 2,496 single nuclei and 54,329 genes, nuclei were kept if the percentage of mapped reads corresponding to ERCC spike-in transcripts was higher than 5 percent but lower than 90 percent. Nuclei needed to have at least 1000 genes detected. Then, genes sequenced in fewer than 25 cells (about 1% of the population) and with read count below 250 reads were filtered out. Known doublets and empty wells from FACS sorting were removed. Finally, only the nuclei having less than 7,000 genes detected and a library size of 10,000 to 300,000 reads were kept. This processing yielded a matrix of 2,016 single nuclei times 19,340 genes.

### ERCC size factor calculation and normalisation

Two different ERCC dilutions were used (1:100,000 and 1:300,000 respectively). When sequencing to saturation, the proportional amounts of sequenced endogenous transcripts and artificial spike-ins depend on the input number of endogenous transcripts. The first step in this normalisation approach is to calculate ERCC size factors as the sum of ERCC reads per nucleus divided by the mean ERCC reads across all nuclei within one dilution. Subsequently, to normalise the expression matrix, the read counts were first divided by the corresponding gene length in kilobases. Then, the cell coverage was corrected by the following approach: counts were summed per nucleus, and the sum was divided by 10,000 times the ERCC size factor, yielding the normalization factor per nucleus. Finally, the reads per nucleus were divided by this factor. Thereby, reads stemming from nuclei with few endogenous reads and many ERCC reads are divided by a smaller factor than reads stemming from nuclei with many endogenous reads and proportionally few ERCC reads.

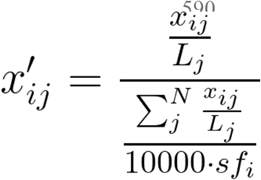

where *x^’^_ij_* is the normalized count of gene *j* in cell *i*, *L_j_* is the length of gene *j* and *sf_i_* is the ERCC size factor of cell *i.*

Additionally, cells with more than 50,000 gene length-normalised total counts were removed. In order to prevent bias in the downstream results, the technical replicates SNI-234(R2) and SNI-235(R2) that contain the same cells as SNI-160 and SNI-116, respectively, were not further considered. This resulted in a final matrix of 1,649 nuclei times 19,258 genes.

The normalised counts were log-transformed by applying a natural logarithm to one plus the values in the count matrix. Batch effects were corrected for using *combat* [122].

### Data visualisation and clustering

The high dimensionality of the normalised expression matrix was reduced by calculating the first 50 principal components. Based on these, a neighborhood graph was constructed based on 15 nearest neighbors, where the connectivities between data points are calculated by Uniform Manifold Approximation and Projection for Dimension Reduction (UMAP) [123]. For the purpose of visualisation, data points were embedded using t-distributed stochastic neighborhood embedding (tSNE) based on the first 15 principal components and with a perplexity of 30.

*Louvain* clustering on the neighbourhood graph with a resolution of 0.2 was used to separate two clusters of non-parenchymal cells from one cluster of hepatocytes [124]. Moreover, *Louvain* clustering with a resolution of 1.2 on the non-parenchymal cluster revealed 7 subclusters. Overlap of marker genes with differential expressed genes for these clusters as well as visual investigation of known markers on the t-sne embedding was used to annotate these subclusters. Known marker genes were adapted from Aizarini et at. [54].

### Cell cycle analysis and inter-cell type correlation

Cell cycle states were established using *cyclone* [57], as implemented in the scran R package [125]. Cells were classified into G1, S and G2M phase, based on their normalized gene counts (Fig. 2B and Supp. Fig. S2). To calculate correlation between cell types, hierarchical clustering was done based on principal components by calculating pearson correlations.

### Differential expression analysis

In order to find differential expressed genes between groups of interest, Welch’s t-tests were done as implemented in the *scanpy* function *rank_genes_groups* with the parameters *method=“t-test”* and *n_genes=19258.* First, this was done between cell types, considering 2n and 4n hepatocytes together as one group, and then, only hepatocytes were taken in order to compare 2n to 4n. Genes that had a log2 fold change of greater than 0.5 and a Bonferroni-adjusted p-value below 0.05 were considered significantly upregulated; genes with a log2 fold change smaller than –0.5 and a Bonferroni-adjusted p-value below 0.05 were considered significantly downregulated.

For visualisation purposes, 40 nuclei were randomly selected per cell type and their respective top 12 differential expressed genes (based on their score calculated by *rank_genes_groups*) were taken to create a heatmap (Fig 2A) using the *ComplexHeatmap* package in R. For visualization purposes, genes with mean expression between 0.1 and 100 were selected in the MA plot (Fig. 3C).

### Gene set enrichment analysis

To find and visualise genes enriched in functionally relevant groups, the python package *gprofiler* was used, focusing on the ontology *biological process (BP).*

### Transcriptional variability

According to the relationship between mean and coefficient of variation, lowly expressed genes have higher transcriptional variability. Hence, before calculating transcriptional variability between groups of interest, genes with mean normalised log-transformed expression smaller than 0.25 were removed, resulting in 2,102 remaining genes. The coefficient of variation (CV) was calculated on the normalised, log-transformed and batch-corrected matrix by making use of the formula described in Canchola *et al.* [126].

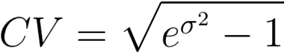

where sigma^2^ is the variation of gene j in the group of interest.

To detect whether there are significant changes in variability between any of the cell types, a Kruskal Wallis test was done and pairwise MannWhitneyU tests afterwards.

Furthermore, highly variable genes per group of interest were identified as genes that have a CV of greater than 1.7. This threshold was chosen after normalising the ERCC-specific reads using the same formula as for the endogenous transcript (with the only adaptation of using 1,000 instead of 10,000 for scaling), log-transforming them, and calculating CV per ERCC, which resulted in 1.66 as the median plus one standard deviation CV for ERCC reads.

### Pseudotemporal ordering based on markers of liver zonation

Non-parenchymal nuclei were removed from the matrix and zonation markers were obtained from [10, 54]. The hereby obtained normalised expression matrix was subset to only contain the zonation marker genes, resulting in a matrix of 1,061 nuclei times 1,742 zonation marker genes. Principle components, k-nearest neighbors and t-SNE as well as UMAP embedding were re-calculated for this matrix as described above. Furthermore, dimensionality reduction was done using diffusion map as described in [106], thus ordering the nuclei along the zonation gradients. Based on marker gene expression and *Louvain* clustering (resolution=1 combining clusters 0, 2 and 3 on the one side, and 1 and 4 on the other), the nuclei were separated into two groups, containing rather pericentral or rather periportal positioned nuclei, respectively. Per group, the percentage of diploid and tetraploid nuclei was calculated. Hereby, a 1.3-time relative enrichment of tetraploid nuclei was observed in the pericentral cluster. For the purpose of better visualisation in the bar plot, the 4n population was subsampled to have the same number of nuclei as the 2n population (Fig 5B).

To visualize changes in expression from pericentral to periportal, the top 30 most differential genes per group were taken and used to construct a PAGA path between these two groups[109]. Furthermore, to visualise the gene expression gradient for individual genes along pseudospace, the vector of diffusion pseudospace was binned into 10 bins between 0 and 1 and mean expression per bin was plotted.

### Co-expression analysis of liver stem cell markers

For this analysis, only hepatocytes were used. A heatmap of known hepatic stem cell markers showed expression of these markers not only in diploid but also in tetraploid nuclei. To investigate whether these markers are expressed in the same nuclei, the matrix was binarized after log-transformation and stored as an additional layer in the AnnData object. This binary matrix was subset to only contain the stem cell markers. Nuclei not expressing any of the stem cell markers were removed, resulting in a matrix of 364 nuclei times 11 stem cell markers. Based on this, pairwise Jaccard distance between the stem cell markers was calculated.

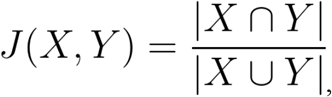

where X is the binary expression vector of gene j_1_ across the nuclei, and Y is the binary expression vector of gene j_2_ across the nuclei.

Furthermore, pairwise Jaccard distances between nuclei and genes, respectively, were used to calculate linkage for hierarchical clustering. By this approach, the gene’s expression level per nucleus is neglected and only its presence or absence is counted.

### Comparison to publicly available single cell RNA-seq data sets

Data were obtained from GEO (Accession numbers GSE84498, GSE124395 [12, 54] and from https://doi.org/10.6084/m9.figshare.5829687.v7 and https://doi.org/10.6084/m9.figshare.5968960.v2 [127]. From unfiltered data, genes with zero expression across cells were removed as well as cells that did not express any genes. Numbers of genes expressed per cell and data set were visualized in violin plots (Fig. 1B).

In order to correlate gene expression between snRNA-seq2 data and *Tabula muris* smart-seq2 data [127], the raw count matrix based on exonic reads was taken. Furthermore, both data sets were subset to only contain hepatocytes. Correlation between log_2_ of the mean gene expression in both datasets was calculated by pearson correlation (Supp. Fig. S1C).

## Abbreviations

CV: Coefficient of Variation
Dpt: diffusion pseudotime
Fig.: Figure
HVG: Highly Variable Genes
DE: Differentially Expressed
DEGs: Differentially Expressed Genes
Supp.: Supplementary
t-SNE: T-distributed stochastic neighbor embedding
UMAP: Uniform Manifold Approximation and Projection for Dimension Reduction

## Supplementary information

Supp. Figures S1

Supp. Figures S2

Supp. Figures S3

Supp. Figures S4

Supp. Figures S5

Supp. Figures S6

Supp. Figures S7

Supp. Table1- Cell type markers (.csv)

Supp. Table2- Gene expression list by cell type (.csv)

Supp. Table3- Gene expression analysis 2n *versus* 4n (.csv)

Supp. Table4- HVG and Coefficient of Variation in 2n and 4n (.csv)

Supp. Table5- Changes in expression distribution in 2n and 4n (.csv)

Supp. Table6- Zonation and DE genes with bins (.csv)

Supp. Table7 – Calculation Sheet for Mosquito HV (.xlsx)

Nextera XT Mosquito HV protocol (pdf)

Nextera XT Mosquito HV protocol (.protocol)

snRNAseq-2 Mosquito HV protocol (pdf)

snRNAseq-2 Mosquito HV protocol (.protocol)

## Acknowledgments

We thank the CRUK-CI core facilities, including Genomics, Flow Cytometry (J. Markovic-Djuric, M. Strzelecki and R. Grenfell) for technical assistance and the Biological Resources Unit (A. Mowbray, N. Jacobs, M. Clayton, and M. Mitchell) for animal husbandry. We are grateful to Dr. M. Hartman, Dr. A. Schröder and Ms. A. Barden (Helmholtz Pioneer Campus) for their legal, managerial and administrative support. We thank Dr. N. Eling for initial data analysis, and Dr. R. Kamies for her comments on the manuscript. We are grateful to Dr. D. T. Odom (CRUK, DKFZ) for his support and scientific advice.

## Authors Contributions

M.L.R, I.K.D., M.C-T. and C.P.M-J., designed experiments; I.D. and C.P.M-J. performed experiments; M.L.R. performed computational analysis. M.L.R, I.K.D, M.C-T and C.P.M-J interpreted the data; E.Ll., and P.C. provided technical assistance; A.D. and C.V. provided computational assistance and advise; C.P.M-J. wrote the manuscript with input from all co-authors. All authors commented on and approved the manuscript.

## Ethical approval and consent to participate

This investigation was approved by the Animal Welfare and Ethics Review Board and followed the Cambridge Institute guidelines for the use of animals in experimental studies under Home Office licences PPL 70/7535 until February 2018 and PPL P9855D13B from March 2018. All animal experimentation was carried out in accordance with the Animals (Scientific Procedures) Act 1986 (United Kingdom) and conformed to the Animal Research: Reporting of In Vivo Experiments (ARRIVE) guidelines developed by the National Centre for the Replacement, Refinement and Reduction of Animals in research (NC3Rs).

---

**Supp. Figure S1.**
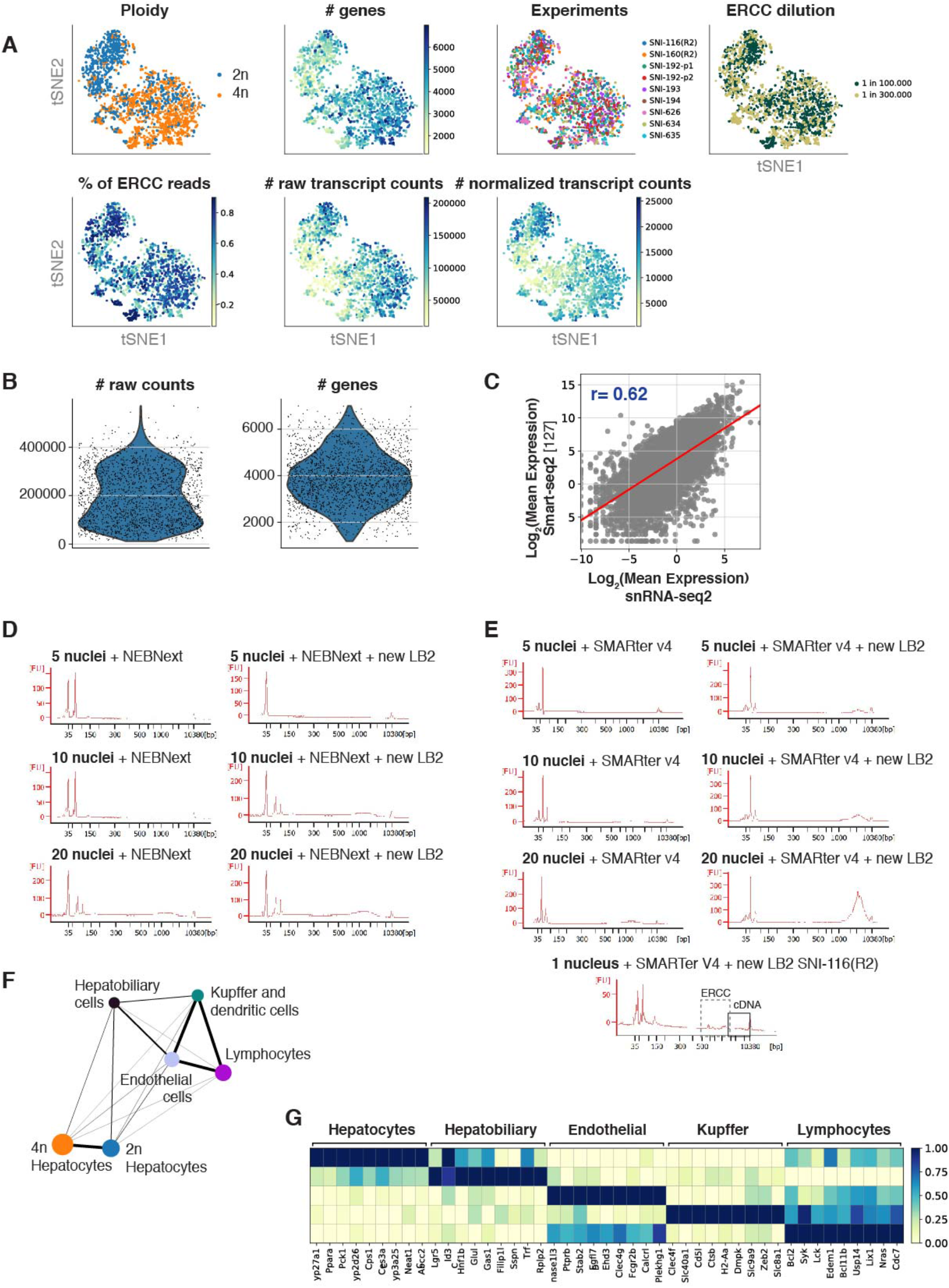
**A)** t-SNE colored by (from left to right) ploidy status, number of genes detected per nucleus, experimental batches, ERCC dilutions, percentage of reads mapping to ERCCs per nucleus, number of raw transcript reads per nucleus, and number of normalized transcript reads per nucleus. **B)** Violin plot showing the distributions of raw counts (left) and number of genes (right). **C)** Scatter plot depicting the mean expression of genes for smart-seq2 single cell data (y-axis) against snRNAseq2 (x-axis), including regression line (rho=0.62). **D)** Bioanalyzer plots showing the increase in cDNA concentration with the new snRNA-seq2 lysis buffer with NEBnext and **E)** with SMARTer v4 Takara. **F)** PAGA showing the connection between cell types. **G)** Matrixplot showing the relative gene expression levels of gene markers of the hepatic cell types identified.

**Supp. Figure S2.**
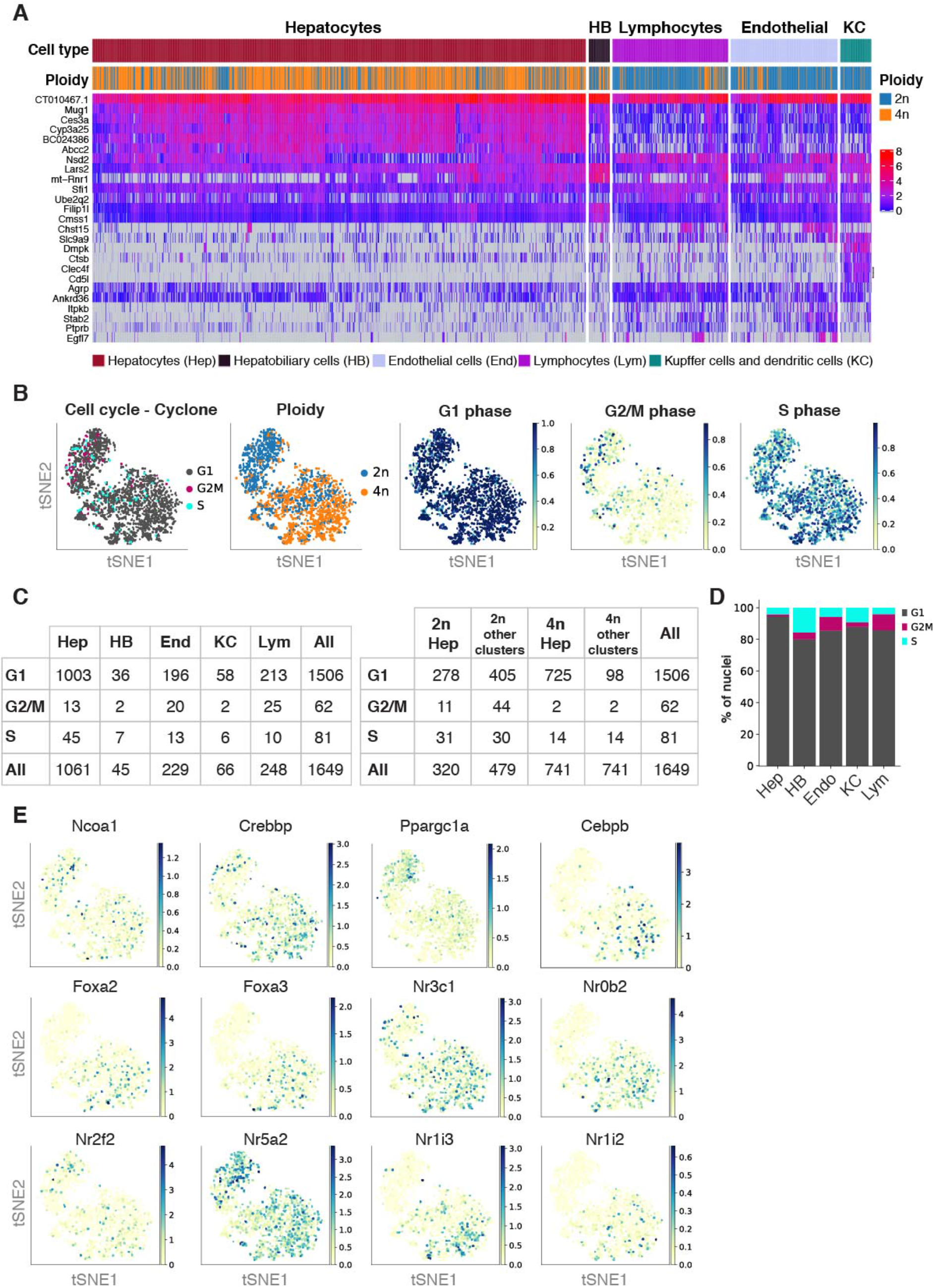
**A)** Heatmap showing the gene expression of the top five differentially expressed genes between cell types. **B)** t-SNE colored by cell cycle phases assigned by *Cyclone,* ploidy status, G1-scores, G2M-scores, and S-scores. **C)** Tables with the number of nuclei in the respective cell cycle phase per cell types (left) and per ploidy level in low resolution *Louvain* clusters (right). **D)** Barplot showing the percentage of nuclei in respective cell cycle phase per cell type. **E)** t-SNE colored by expression level of key transcription factors

**Supp. Figure S3.**
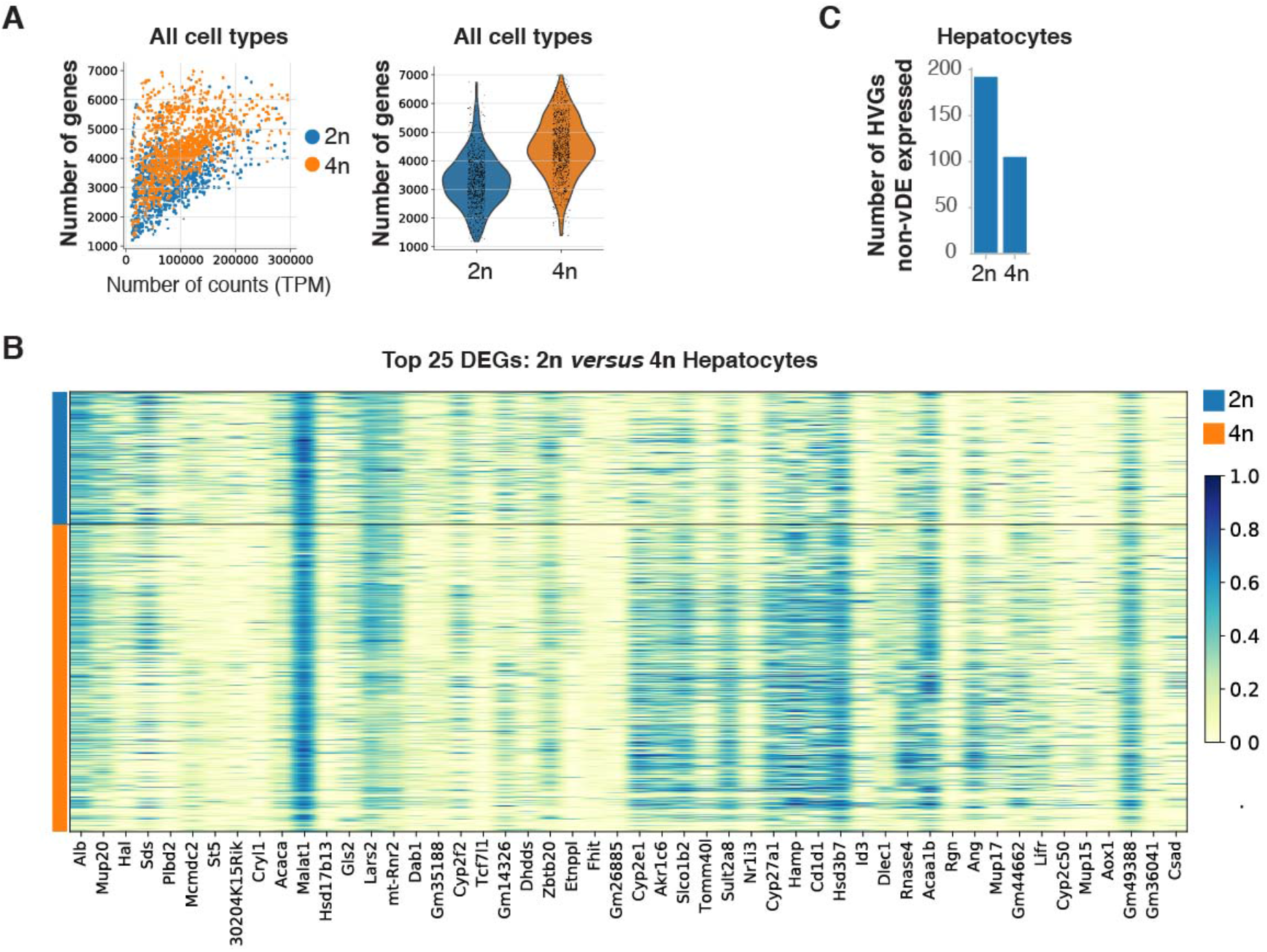
**A)** Scatterplot showing the number of genes per normalized counts for all cells colored by their ploidy level (left); Violin plot showing a 1.35-times increase in number of genes detected from 4n nuclei in comparison to 2n nuclei (right). **B)** Heatmap showing gene expression of the top 25 differentially expressed genes between 2n and 4n hepatocytes. **C)** Bar plot showing the number of highly variable genes (HVG) that are not differentially expressed between 2n and 4n hepatocytes

**Supp. Figure S4.**
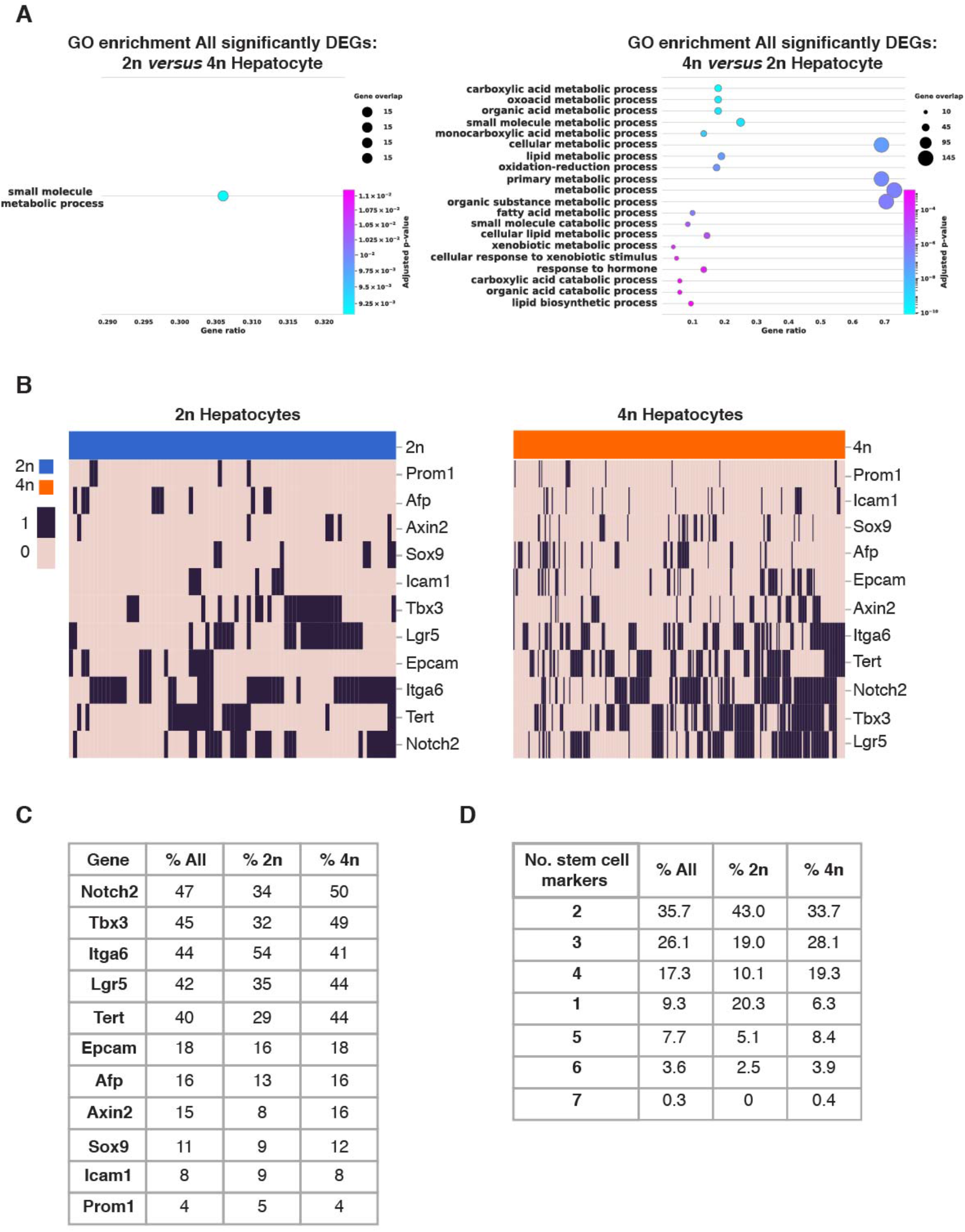
**A)** Gene set enrichment plots of all differentially expressed genes in 2n hepatocytes (left), 4n hepatocytes (right). **B)** Heatmap showing binary expression of stem cell markers in 2n (left) and 4n hepatocytes (right); linkage is calculated based on Jaccard distances. **C)** Percentage of cell in which stem/progenitor markers are detected in all, in 2n hepatocytes and 4n hepatocytes. **D)** Percentage of cells in which stem cell markers that are co-expressed in all, in 2n hepatocytes and 4n hepatocytes.

**Supp. Figure S5.**
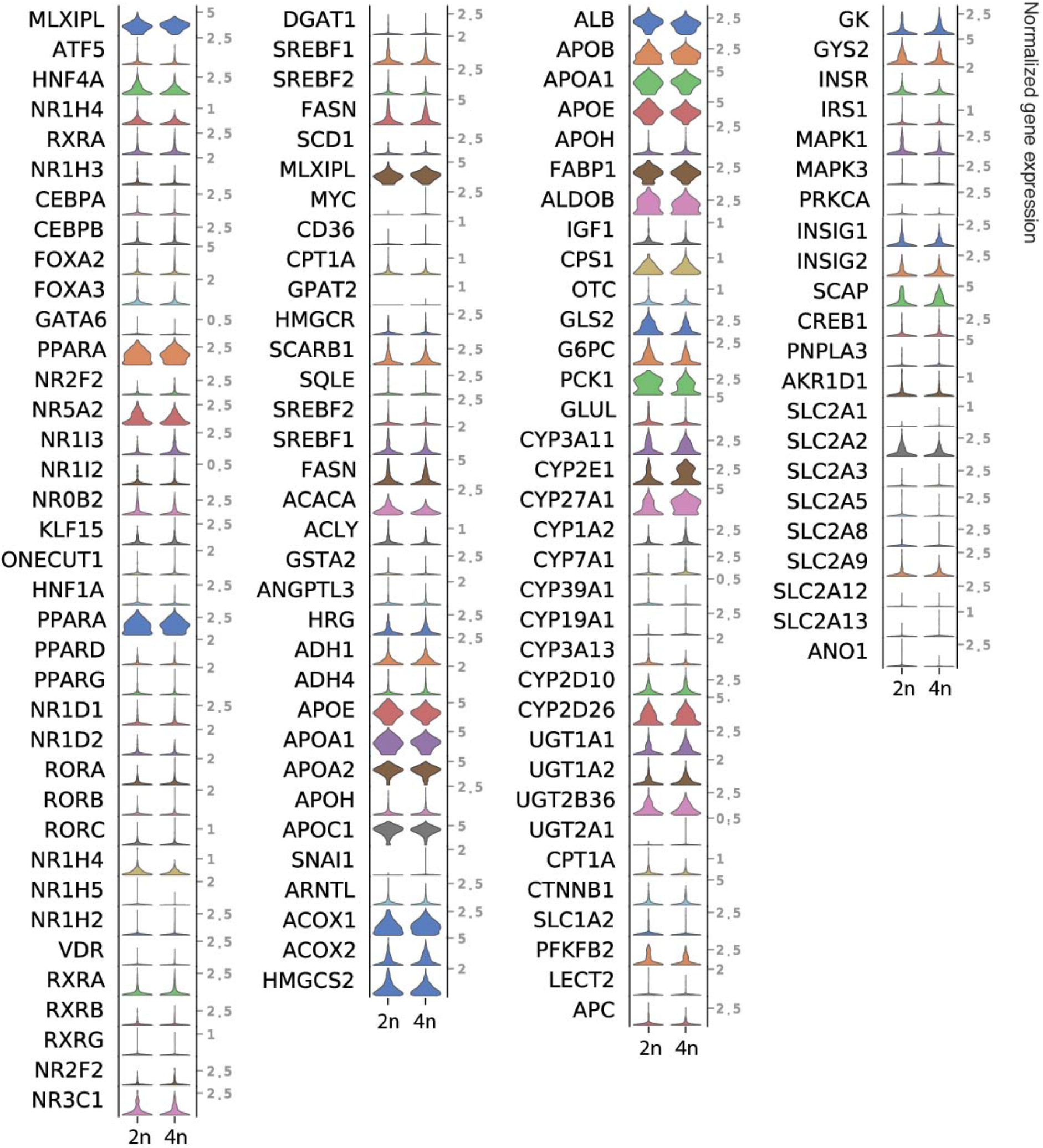
Violin plots showing the expression profiles of relevant genes in hepatic energy homeostasis and metabolism of xenobiotics.

**Supp. Figure S6.**
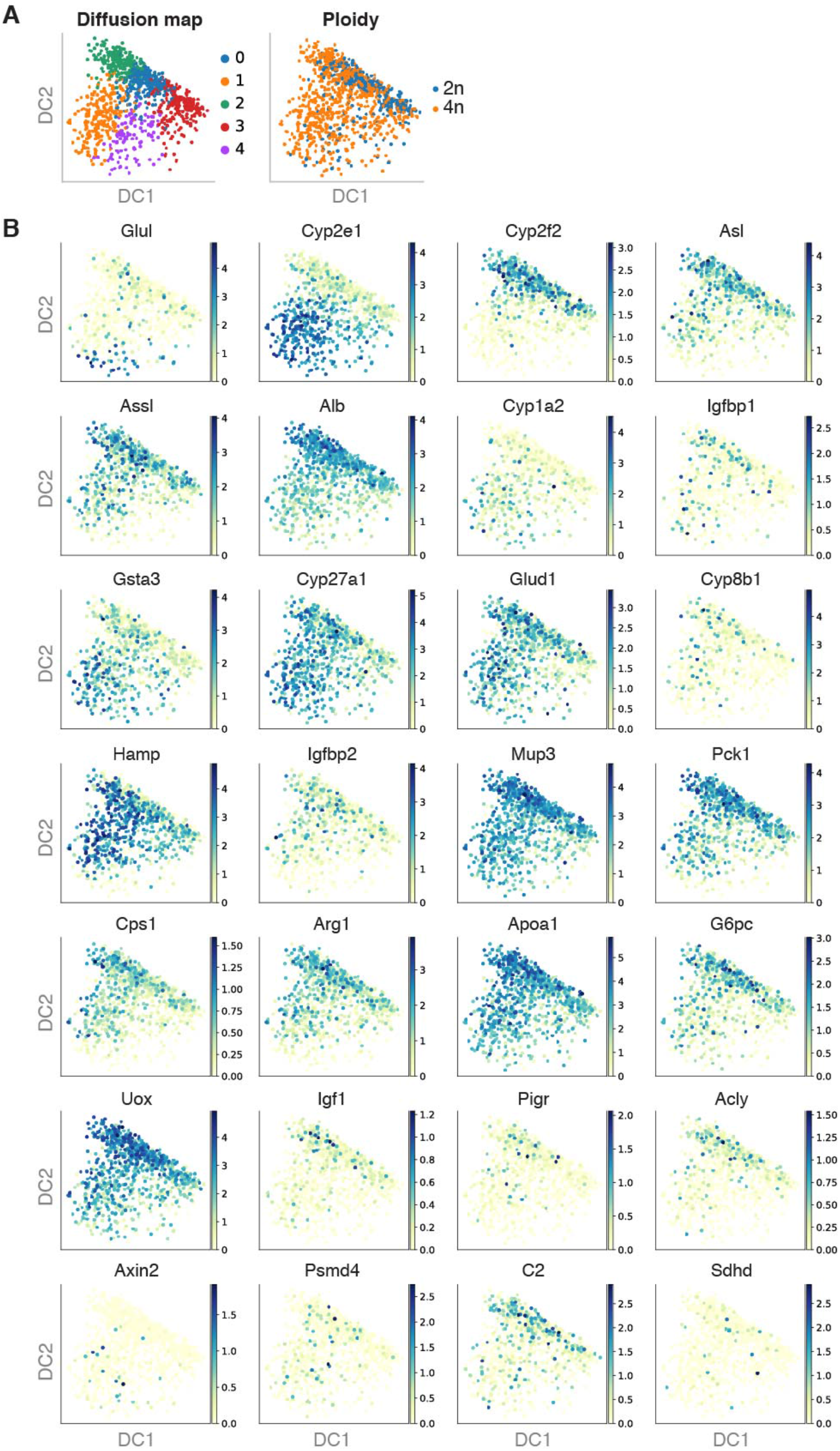
**A** Diffusion map colored by *Louvain* clustering (left) and ploidy (right). **B** Diffusion maps colored by zonation markers described in Halpern *et al.* 2017 [12]

**Supp. Figure S7.**
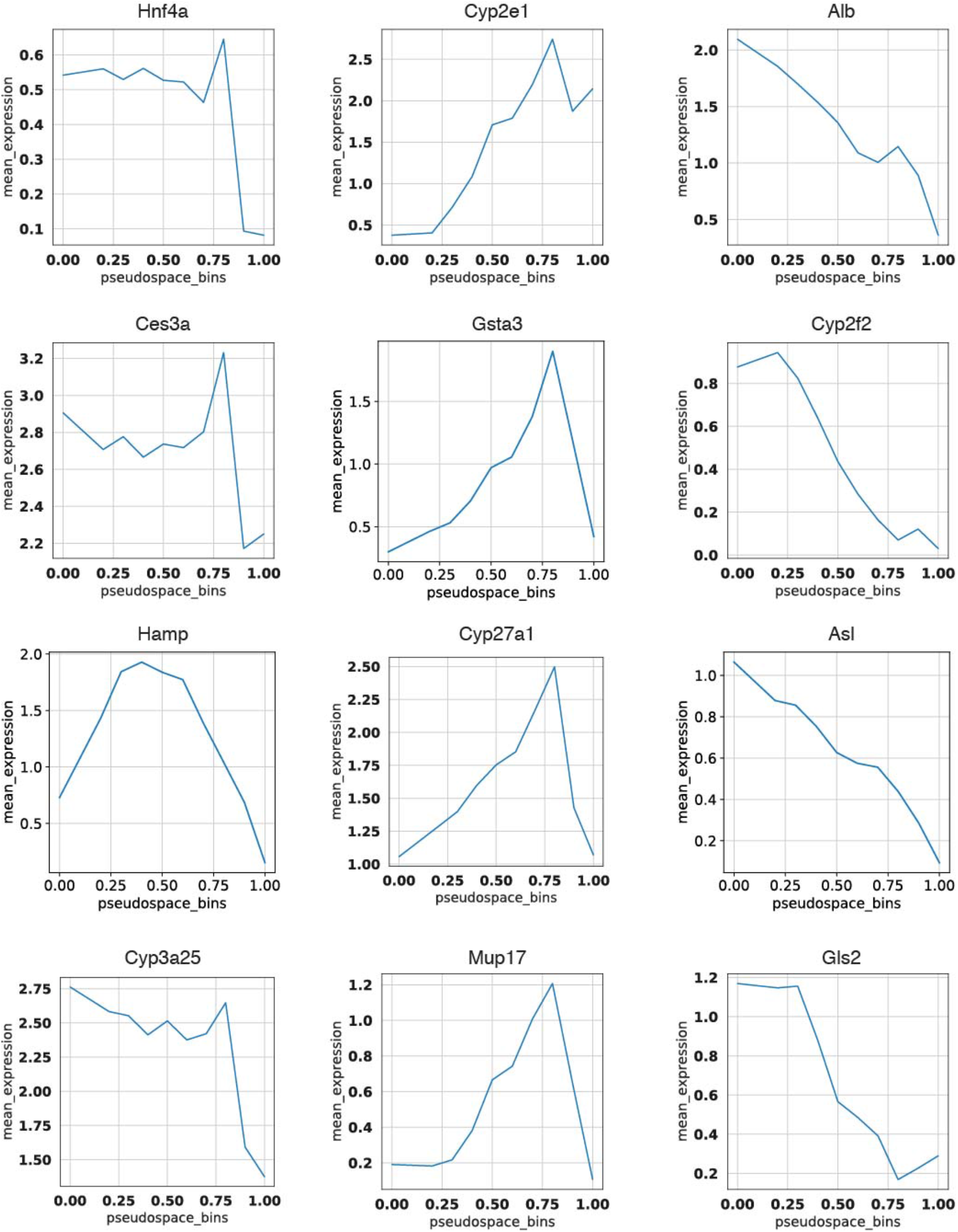
Line plots depicting mean expression of individual zonation markers per bin of the diffusion pseudospace vector.

## References

1. Martinez-Jimenez CP, Castell JV, Gomez-Lechon MJ, Jover R: Transcriptional activation of CYP2C9, CYP1A1, and CYP1A2 by hepatocyte nuclear factor 4alpha requires coactivators peroxisomal proliferator activated receptor-gamma coactivator 1alpha and steroid receptor coactivator 1. Mol Pharmacol 2006, 70:1681–1692.

2. Martinez-Jimenez CP, Gomez-Lechon MJ, Castell JV, Jover R: Underexpressed coactivators PGC1alpha and SRC1 impair hepatocyte nuclear factor 4 alpha function and promote dedifferentiation in human hepatoma cells. J Biol Chem 2006, 281:29840–29849.

3. Martinez-Jimenez CP, Jover R, Donato MT, Castell JV, Gomez-Lechon MJ: Transcriptional regulation and expression of CYP3A4 in hepatocytes. Curr Drug Metab 2007, 8:185-194.

4. Castell JV, Jover R, Martinez-Jimenez CP, Gomez-Lechon MJ: Hepatocyte cell lines: their use, scope and limitations in drug metabolism studies. Expert Opin Drug Metab Toxicol 2006, 2:183–212.

5. Luecken MD, Theis FJ: Current best practices in single-cell RNA-seq analysis: a tutorial. Mol Syst Biol 2019, 15:e8746.

6. Martinez-Jimenez CP, Kyrmizi I, Cardot P, Gonzalez FJ, Talianidis I: Hepatocyte nuclear factor 4alpha coordinates a transcription factor network regulating hepatic fatty acid metabolism. Mol Cell Biol 2010, 30:565–577.

7. Schmidt D, Wilson MD, Ballester B, Schwalie PC, Brown GD, Marshall A, Kutter C, Watt S, Martinez-Jimenez CP, Mackay S, et al: Five-vertebrate ChIP-seq reveals the evolutionary dynamics of transcription factor binding. Science 2010, 328:1036–1040.

8. Morales-Navarrete H, Segovia-Miranda F, Klukowski P, Meyer K, Nonaka H, Marsico G, Chernykh M, Kalaidzidis A, Zerial M, Kalaidzidis Y: A versatile pipeline for the multi-scale digital reconstruction and quantitative analysis of 3D tissue architecture. Elife 2015, 4.

9. Ben-Moshe S, Itzkovitz S: Spatial heterogeneity in the mammalian liver. Nat Rev Gastroenterol Hepatol 2019, 16:395–410.

10. Ben-Moshe S, Shapira Y, Moor AE, Manco R, Veg T, Bahar Halpern K, Itzkovitz S: Spatial sorting enables comprehensive characterization of liver zonation. Nat Metab 2019, 1:899–911.

11. Halpern KB, Shenhav R, Massalha H, Toth B, Egozi A, Massasa EE, Medgalia C, David E, Giladi A, Moor AE, et al: Paired-cell sequencing enables spatial gene expression mapping of liver endothelial cells. Nat Biotechnol 2018, 36:962–970.

12. Halpern KB, Shenhav R, Matcovitch-Natan O, Toth B, Lemze D, Golan M, Massasa EE, Baydatch S, Landen S, Moor AE, et al: Single-cell spatial reconstruction reveals global division of labour in the mammalian liver. Nature 2017, 542:352–356.

13. Moor AE, Itzkovitz S: Spatial transcriptomics: paving the way for tissue-level systems biology. Curr Opin Biotechnol 2017, 46:126–133.

14. Dobie R, Wilson-Kanamori JR, Henderson BEP, Smith JR, Matchett KP, Portman JR, Wallenborg K, Picelli S, Zagorska A, Pendem SV, et al: Single-Cell Transcriptomics Uncovers Zonation of Function in the Mesenchyme during Liver Fibrosis. Cell Rep 2019, 29:1832–1847 e1838.

15. Ramachandran P, Dobie R, Wilson-Kanamori JR, Dora EF, Henderson BEP, Luu NT, Portman JR, Matchett KP, Brice M, Marwick JA, et al: Resolving the fibrotic niche of human liver cirrhosis at single-cell level. Nature 2019, 575:512–518.

16. Ramachandran P, Matchett KP, Dobie R, Wilson-Kanamori JR, Henderson NC: Single-cell technologies in hepatology: new insights into liver biology and disease pathogenesis. Nature Reviews Gastroenterology & Hepatology 2020.

17. van den Brink SC, Sage F, Vertesy A, Spanjaard B, Peterson-Maduro J, Baron CS, Robin C, van Oudenaarden A: Single-cell sequencing reveals dissociation-induced gene expression in tissue subpopulations. Nat Methods 2017, 14:935–936.

18. Gomez-Lechon MJ, Donato MT, Castell JV, Jover R: Human hepatocytes in primary culture: the choice to investigate drug metabolism in man. Curr Drug Metab 2004, 5:443–462.

19. Gomez-Lechon MJ, Donato MT, Castell JV, Jover R: Human hepatocytes as a tool for studying toxicity and drug metabolism. Curr Drug Metab 2003, 4:292–312.

20. Grindberg RV, Yee-Greenbaum JL, McConnell MJ, Novotny M, O’Shaughnessy AL, Lambert GM, Arauzo-Bravo MJ, Lee J, Fishman M, Robbins GE, et al: RNA-sequencing from single nuclei. Proc Natl Acad Sci U S A 2013, 110:19802–19807.

21. Krishnaswami SR, Grindberg RV, Novotny M, Venepally P, Lacar B, Bhutani K, Linker SB, Pham S, Erwin JA, Miller JA, et al: Using single nuclei for RNA-seq to capture the transcriptome of postmortem neurons. Nat Protoc 2016, 11:499–524.

22. Kalish BT, Barkat TR, Diel EE, Zhang EJ, Greenberg ME, Hensch TK: Single-nucleus RNA sequencing of mouse auditory cortex reveals critical period triggers and brakes. Proc Natl Acad Sci U S A 2020, 117:11744–11752.

23. Lake BB, Codeluppi S, Yung YC, Gao D, Chun J, Kharchenko PV, Linnarsson S, Zhang K: A comparative strategy for single-nucleus and single-cell transcriptomes confirms accuracy in predicted cell-type expression from nuclear RNA. Sci Rep 2017, 7:6031.

24. Lake BB, Ai R, Kaeser GE, Salathia NS, Yung YC, Liu R, Wildberg A, Gao D, Fung HL, Chen S, et al: Neuronal subtypes and diversity revealed by single-nucleus RNA sequencing of the human brain. Science 2016, 352:1586–1590.

25. Habib N, Li Y, Heidenreich M, Swiech L, Avraham-Davidi I, Trombetta JJ, Hession C, Zhang F, Regev A: Div-Seq: Single-nucleus RNA-Seq reveals dynamics of rare adult newborn neurons. Science 2016, 353:925–928.

26. Habib N., Basu A., Avraham-Davidi I., Burks T., Choudhury SR., Aguet F., Gelfand E., Ardlie K., Weitz DA., Rozenblatt-Rosen O., et al: DroNc-Seq: Deciphering cell types in human archived brain tissues by massively-parallel single nucleus RNA-seq. bioRxiv 2017, 9:1–12.

27. Hu P, Fabyanic E, Kwon DY, Tang S, Zhou Z, Wu H: Dissecting Cell-Type Composition and Activity-Dependent Transcriptional State in Mammalian Brains by Massively Parallel Single-Nucleus RNA-Seq. Mol Cell 2017, 68:1006–1015 e1007.

28. Koenitzer JR, Wu H, Atkinson JJ, Brody SL, Humphreys BD: Single nucleus RNASeq profiling of mouse lung: reduced dissociation bias and improved detection of rare cell types compared with single cell RNASeq. bioRxiv 2020:2020.2003.2006.981407.

29. Wu H, Kirita Y, Donnelly EL, Humphreys BD: Advantages of Single-Nucleus over Single-Cell RNA Sequencing of Adult Kidney: Rare Cell Types and Novel Cell States Revealed in Fibrosis. J Am Soc Nephrol 2019, 30:23–32.

30. Wilson PC, Wu H, Kirita Y, Uchimura K, Ledru N, Rennke HG, Welling PA, Waikar SS, Humphreys BD: The single-cell transcriptomic landscape of early human diabetic nephropathy. Proc Natl Acad Sci U S A 2019, 116:19619–19625.

31. Denisenko E, Guo BB, Jones M, Hou R, de Kock L, Lassmann T, Poppe D, Clement O, Simmons RK, Lister R, Forrest ARR: Systematic assessment of tissue dissociation and storage biases in single-cell and single-nucleus RNA-seq workflows. Genome Biol 2020, 21:130.

32. Lake BB, Chen S, Hoshi M, Plongthongkum N, Salamon D, Knoten A, Vijayan A, Venkatesh R, Kim EH, Gao D, et al: A single-nucleus RNA-sequencing pipeline to decipher the molecular anatomy and pathophysiology of human kidneys. Nat Commun 2019, 10:2832.

33. Wolfien M, Galow AM, Muller P, Bartsch M, Brunner RM, Goldammer T, Wolkenhauer O, Hoeflich A, David R: Single-Nucleus Sequencing of an Entire Mammalian Heart: Cell Type Composition and Velocity. Cells 2020, 9.

34. Selewa A, Dohn R, Eckart H, Lozano S, Xie B, Gauchat E, Elorbany R, Rhodes K, Burnett J, Gilad Y, et al: Systematic Comparison of High-throughput Single-Cell and Single-Nucleus Transcriptomes during Cardiomyocyte Differentiation. Sci Rep 2020, 10:1535.

35. Ding J, Adiconis X, Simmons SK, Kowalczyk MS, Hession CC, Marjanovic ND, Hughes TK, Wadsworth MH, Burks T, Nguyen LT, et al: Systematic comparison of single-cell and single-nucleus RNA-sequencing methods. Nat Biotechnol 2020.

36. Slyper M, Porter CBM, Ashenberg O, Waldman J, Drokhlyansky E, Wakiro I, Smillie C, Smith-Rosario G, Wu J, Dionne D, et al: A single-cell and single-nucleus RNA-Seq toolbox for fresh and frozen human tumors. Nat Med 2020, 26:792–802.

37. Abdelmoez MN, Iida K, Oguchi Y, Nishikii H, Yokokawa R, Kotera H, Uemura S, Santiago JG, Shintaku H: SINC-seq: correlation of transient gene expressions between nucleus and cytoplasm reflects single-cell physiology. Genome Biol 2018, 19:66.

38. Duncan AW, Taylor MH, Hickey RD, Hanlon Newell AE, Lenzi ML, Olson SB, Finegold MJ, Grompe M: The ploidy conveyor of mature hepatocytes as a source of genetic variation. Nature 2010, 467:707–710.

39. Duncan AW, Hanlon Newell AE, Smith L, Wilson EM, Olson SB, Thayer MJ, Strom SC, Grompe M: Frequent aneuploidy among normal human hepatocytes. Gastroenterology 2012, 142:25–28.

40. Donne R, Saroul-Ainama M, Cordier P, Celton-Morizur S, Desdouets C: Polyploidy in liver development, homeostasis and disease. Nat Rev Gastroenterol Hepatol 2020.

41. Epstein CJ: Cell size, nuclear content, and the development of polyploidy in the Mammalian liver. Proc Natl Acad Sci U S A 1967, 57:327–334.

42. Kreutz C, MacNelly S, Follo M, Waldin A, Binninger-Lacour P, Timmer J, Bartolome-Rodriguez MM: Hepatocyte Ploidy Is a Diversity Factor for Liver Homeostasis. Front Physiol 2017, 8:862.

43. Lu P, Prost S, Caldwell H, Tugwood JD, Betton GR, Harrison DJ: Microarray analysis of gene expression of mouse hepatocytes of different ploidy. Mamm Genome 2007, 18:617–626.

44. Buettner F, Natarajan KN, Casale FP, Proserpio V, Scialdone A, Theis FJ, Teichmann SA, Marioni JC, Stegle O: Computational analysis of cell-to-cell heterogeneity in single-cell RNA-sequencing data reveals hidden subpopulations of cells. Nat Biotechnol 2015, 33:155–160.

45. Lun ATL, Marioni JC: Overcoming confounding plate effects in differential expression analyses of single-cell RNA-seq data. Biostatistics 2017, 18:451–464.

46. Martinez-Jimenez CP, Eling N, Chen HC, Vallejos CA, Kolodziejczyk AA, Connor F, Stojic L, Rayner TF, Stubbington MJ, Teichmann SA, et al: Aging increases cell-to-cell transcriptional variability upon immune stimulation. Science 2017, 355:1433–1436.

47. Lun ATL, Calero-Nieto FJ, Haim-Vilmovsky L, Gottgens B, Marioni JC: Assessing the reliability of spike-in normalization for analyses of single-cell RNA sequencing data. Genome Res 2017, 27:1795–1806.

48. Eling N, Morgan MD, Marioni JC: Challenges in measuring and understanding biological noise. Nat Rev Genet 2019, 20:536–548.

49. Svensson V, Natarajan KN, Ly LH, Miragaia RJ, Labalette C, Macaulay IC, Cvejic A, Teichmann SA: Power analysis of single-cell RNA-sequencing experiments. Nat Methods 2017, 14:381–387.

50. Sathyamurthy A, Johnson KR, Matson KJE, Dobrott CI, Li L, Ryba AR, Bergman TB, Kelly MC, Kelley MW, Levine AJ: Massively Parallel Single Nucleus Transcriptional Profiling Defines Spinal Cord Neurons and Their Activity during Behavior. Cell Rep 2018, 22:2216–2225.

51. Gao R, Kim C, Sei E, Foukakis T, Crosetto N, Chan LK, Srinivasan M, Zhang H, Meric-Bernstam F, Navin N: Nanogrid single-nucleus RNA sequencing reveals phenotypic diversity in breast cancer. Nat Commun 2017, 8:228.

52. Mereu E, Lafzi A, Moutinho C, Ziegenhain C, MacCarthy DJ, Alvarez A, Batlle E, Sagar, Grün D, Lau JK, et al: Benchmarking Single-Cell RNA Sequencing Protocols for Cell Atlas Projects. bioRxiv 2019:630087.

53. Segal JM, Kent D, Wesche DJ, Ng SS, Serra M, Oules B, Kar G, Emerton G, Blackford SJI, Darmanis S, et al: Single cell analysis of human foetal liver captures the transcriptional profile of hepatobiliary hybrid progenitors. Nat Commun 2019, 10:3350.

54. Aizarani N, Saviano A, Sagar, Mailly L, Durand S, Herman JS, Pessaux P, Baumert TF, Grun D: A human liver cell atlas reveals heterogeneity and epithelial progenitors. Nature 2019, 572:199–204.

55. Celton-Morizur S, Desdouets C: Polyploidization of liver cells. Adv Exp Med Biol 2010, 676:123–135.

56. McDavid A, Finak G, Gottardo R: The contribution of cell cycle to heterogeneity in single-cell RNA-seq data. Nat Biotechnol 2016, 34:591–593.

57. Scialdone A, Natarajan KN, Saraiva LR, Proserpio V, Teichmann SA, Stegle O, Marioni JC, Buettner F: Computational assignment of cell-cycle stage from single-cell transcriptome data. Methods 2015, 85:54–61.

58. Martinez-Jimenez CP, Gomez-Lechon MJ, Castell JV, Jover R: Transcriptional regulation of the human hepatic CYP3A4: identification of a new distal enhancer region responsive to CCAAT/enhancer-binding protein beta isoforms (liver activating protein and liver inhibitory protein). Mol Pharmacol 2005, 67:2088–2101.

59. Montagner A, Polizzi A, Fouche E, Ducheix S, Lippi Y, Lasserre F, Barquissau V, Regnier M, Lukowicz C, Benhamed F, et al: Liver PPARalpha is crucial for whole-body fatty acid homeostasis and is protective against NAFLD. Gut 2016, 65:1202–1214.

60. Lee SS, Pineau T, Drago J, Lee EJ, Owens JW, Kroetz DL, Fernandez-Salguero PM, Westphal H, Gonzalez FJ: Targeted disruption of the alpha isoform of the peroxisome proliferator-activated receptor gene in mice results in abolishment of the pleiotropic effects of peroxisome proliferators. Mol Cell Biol 1995, 15:3012–3022.

61. Pawlak M, Lefebvre P, Staels B: Molecular mechanism of PPARalpha action and its impact on lipid metabolism, inflammation and fibrosis in non-alcoholic fatty liver disease. J Hepatol 2015, 62:720–733.

62. Yamashita H, Takenoshita M, Sakurai M, Bruick RK, Henzel WJ, Shillinglaw W, Arnot D, Uyeda K: A glucose-responsive transcription factor that regulates carbohydrate metabolism in the liver. Proc Natl Acad Sci U S A 2001, 98:9116–9121.

63. Benhamed F, Denechaud PD, Lemoine M, Robichon C, Moldes M, Bertrand-Michel J, Ratziu V, Serfaty L, Housset C, Capeau J, et al: The lipogenic transcription factor ChREBP dissociates hepatic steatosis from insulin resistance in mice and humans. J Clin Invest 2012, 122:2176–2194.

64. Samuel VT, Shulman GI: Nonalcoholic Fatty Liver Disease as a Nexus of Metabolic and Hepatic Diseases. Cell Metab 2018, 27:22–41.

65. Casagrande V, Mauriello A, Bischetti S, Mavilio M, Federici M, Menghini R: Hepatocyte specific TIMP3 expression prevents diet dependent fatty liver disease and hepatocellular carcinoma. Sci Rep 2017, 7:6747.

66. Nikolaou N, Gathercole LL, Marchand L, Althari S, Dempster NJ, Green CJ, van de Bunt M, McNeil C, Arvaniti A, Hughes BA, et al: AKR1D1 is a novel regulator of metabolic phenotype in human hepatocytes and is dysregulated in non-alcoholic fatty liver disease. Metabolism 2019, 99:67–80.

67. Xiong X, Kuang H, Ansari S, Liu T, Gong J, Wang S, Zhao XY, Ji Y, Li C, Guo L, et al: Landscape of Intercellular Crosstalk in Healthy and NASH Liver Revealed by Single-Cell Secretome Gene Analysis. Mol Cell 2019, 75:644–660 e645.

68. Wang X, Zheng Z, Caviglia JM, Corey KE, Herfel TM, Cai B, Masia R, Chung RT, Lefkowitch JH, Schwabe RF, Tabas I: Hepatocyte TAZ/WWTR1 Promotes Inflammation and Fibrosis in Nonalcoholic Steatohepatitis. Cell Metab 2016, 24:848–862.

69. van Koppen A, Verschuren L, van den Hoek AM, Verheij J, Morrison MC, Li K, Nagabukuro H, Costessi A, Caspers MPM, van den Broek TJ, et al: Uncovering a Predictive Molecular Signature for the Onset of NASH-Related Fibrosis in a Translational NASH Mouse Model. Cell Mol Gastroenterol Hepatol 2018, 5:83–98 e10.

70. Ramnath D, Irvine KM, Lukowski SW, Horsfall LU, Loh Z, Clouston AD, Patel PJ, Fagan KJ, Iyer A, Lampe G, et al: Hepatic expression profiling identifies steatosis-independent and steatosis-driven advanced fibrosis genes. JCI Insight 2018, 3.

71. MacParland SA, Liu JC, Ma XZ, Innes BT, Bartczak AM, Gage BK, Manuel J, Khuu N, Echeverri J, Linares I, et al: Single cell RNA sequencing of human liver reveals distinct intrahepatic macrophage populations. Nat Commun 2018, 9:4383.

72. Krenkel O, Hundertmark J, Ritz TP, Weiskirchen R, Tacke F: Single Cell RNA Sequencing Identifies Subsets of Hepatic Stellate Cells and Myofibroblasts in Liver Fibrosis. Cells 2019, 8.

73. Pepe-Mooney BJ, Dill MT, Alemany A, Ordovas-Montanes J, Matsushita Y, Rao A, Sen A, Miyazaki M, Anakk S, Dawson PA, et al: Single-Cell Analysis of the Liver Epithelium Reveals Dynamic Heterogeneity and an Essential Role for YAP in Homeostasis and Regeneration. Cell Stem Cell 2019, 25:23–38 e28.

74. Su X, Shi Y, Zou X, Lu ZN, Xie G, Yang JYH, Wu CC, Cui XF, He KY, Luo Q, et al: Single-cell RNA-Seq analysis reveals dynamic trajectories during mouse liver development. BMC Genomics 2017, 18:946.

75. Fox DT, Duronio RJ: Endoreplication and polyploidy: insights into development and disease. Development 2013, 140:3–12.

76. Katsuda T, Hosaka K, Matsuzaki J, Usuba W, Prieto-Vila M, Yamaguchi T, Tsuchiya A, Terai S, Ochiya T: Transcriptomic Dissection of Hepatocyte Heterogeneity: Linking Ploidy, Zonation, and Stem/Progenitor Cell Characteristics. Cell Mol Gastroenterol Hepatol 2020, 9:161–183.

77. Matsumoto T, Wakefield L, Tarlow BD, Grompe M: In Vivo Lineage Tracing of Polyploid Hepatocytes Reveals Extensive Proliferation during Liver Regeneration. Cell Stem Cell 2020, 26:34–47 e33.

78. Davoli T, de Lange T: The causes and consequences of polyploidy in normal development and cancer. Annu Rev Cell Dev Biol 2011, 27:585–610.

79. Wang MJ, Chen F, Lau JTY, Hu YP: Hepatocyte polyploidization and its association with pathophysiological processes. Cell Death Dis 2017, 8:e2805.

80. Gjelsvik KJ, Besen-McNally R, Losick VP: Solving the Polyploid Mystery in Health and Disease. Trends Genet 2019, 35:6–14.

81. Turner R, Lozoya O, Wang Y, Cardinale V, Gaudio E, Alpini G, Mendel G, Wauthier E, Barbier C, Alvaro D, Reid LM: Human hepatic stem cell and maturational liver lineage biology. Hepatology 2011, 53:1035–1045.

82. Schwartz-Arad D, Zajicek G, Bartfeld E: The streaming liver IV: DNA content of the hepatocyte increases with its age. Liver 1989, 9:93–99.

83. Kudryavtsev BN, Kudryavtseva MV, Sakuta GA, Stein GI: Human hepatocyte polyploidization kinetics in the course of life cycle. Virchows Arch B Cell Pathol Incl Mol Pathol 1993, 64:387–393.

84. Gentric G, Maillet V, Paradis V, Couton D, L’Hermitte A, Panasyuk G, Fromenty B, Celton-Morizur S, Desdouets C: Oxidative stress promotes pathologic polyploidization in nonalcoholic fatty liver disease. J Clin Invest 2015, 125:981–992.

85. Bou-Nader M, Caruso S, Donne R, Celton-Morizur S, Calderaro J, Gentric G, Cadoux M, L’Hermitte A, Klein C, Guilbert T, et al: Polyploidy spectrum: a new marker in HCC classification. Gut 2019.

86. Cobbina E, Akhlaghi F: Non-alcoholic fatty liver disease (NAFLD) -pathogenesis, classification, and effect on drug metabolizing enzymes and transporters. Drug Metab Rev 2017, 49:197–211.

87. Rice AM, McLysaght A: Dosage-sensitive genes in evolution and disease. BMC Biol 2017, 15:78.

88. Bahar Halpern K, Tanami S, Landen S, Chapal M, Szlak L, Hutzler A, Nizhberg A, Itzkovitz S: Bursty gene expression in the intact mammalian liver. Mol Cell 2015, 58:147–156.

89. Grun D, Kester L, van Oudenaarden A: Validation of noise models for single-cell transcriptomics. Nat Methods 2014, 11:637–640.

90. Goolam M, Scialdone A, Graham SJL, Macaulay IC, Jedrusik A, Hupalowska A, Voet T, Marioni JC, Zernicka-Goetz M: Heterogeneity in Oct4 and Sox2 Targets Biases Cell Fate in 4-Cell Mouse Embryos. Cell 2016, 165:61–74.

91. Canchola JA, Tang S, Hemyari P, Paxinos E, Marins E: Correct use of percent coefficient of variation (%CV) formula for log-transformed data. MOJ Proteomics & Bioinformatics 2017, 6:316–317.

92. Sun T, Pikiolek M, Orsini V, Bergling S, Holwerda S, Morelli L, Hoppe PS, Planas-Paz L, Yang Y, Ruffner H, et al: AXIN2(+) Pericentral Hepatocytes Have Limited Contributions to Liver Homeostasis and Regeneration. Cell Stem Cell 2020, 26:97–107 e106.

93. Wilkinson PD, Alencastro F, Delgado ER, Leek MP, Weirich MP, Otero PA, Roy N, Brown WK, Oertel M, Duncan AW: Polyploid Hepatocytes Facilitate Adaptation and Regeneration to Chronic Liver Injury. Am J Pathol 2019, 189:1241–1255.

94. Zhang S, Nguyen LH, Zhou K, Tu HC, Sehgal A, Nassour I, Li L, Gopal P, Goodman J, Singal AG, et al: Knockdown of Anillin Actin Binding Protein Blocks Cytokinesis in Hepatocytes and Reduces Liver Tumor Development in Mice Without Affecting Regeneration. Gastroenterology 2018, 154:1421–1434.

95. Zhang S, Zhou K, Luo X, Li L, Tu HC, Sehgal A, Nguyen LH, Zhang Y, Gopal P, Tarlow BD, et al: The Polyploid State Plays a Tumor-Suppressive Role in the Liver. Dev Cell 2018, 44:447–459 e445.

96. Wang B, Zhao L, Fish M, Logan CY, Nusse R: Self-renewing diploid Axin2(+) cells fuel homeostatic renewal of the liver. Nature 2015, 524:180–185.

97. Chen F, Jimenez RJ, Sharma K, Luu HY, Hsu BY, Ravindranathan A, Stohr BA, Willenbring H: Broad Distribution of Hepatocyte Proliferation in Liver Homeostasis and Regeneration. Cell Stem Cell 2020, 26:27–33 e24.

98. Levandowsky M, Winter D: Distance between Sets. Nature 1971, 234:34–35.

99. Fuxman Bass JI, Diallo A, Nelson J, Soto JM, Myers CL, Walhout AJ: Using networks to measure similarity between genes: association index selection. Nat Methods 2013, 10:1169–1176.

100. Li X, Lalic J, Baeza-Centurion P, Dhar R, Lehner B: Changes in gene expression predictably shift and switch genetic interactions. Nat Commun 2019, 10:3886.

101. Shalek AK, Satija R, Adiconis X, Gertner RS, Gaublomme JT, Raychowdhury R, Schwartz S, Yosef N, Malboeuf C, Lu D, et al: Single-cell transcriptomics reveals bimodality in expression and splicing in immune cells. Nature 2013, 498:236–240.

102. Chalancon G, Ravarani CN, Balaji S, Martinez-Arias A, Aravind L, Jothi R, Babu MM: Interplay between gene expression noise and regulatory network architecture. Trends Genet 2012, 28:221–232.

103. Anatskaya OV, Vinogradov AE: Genome multiplication as adaptation to tissue survival: evidence from gene expression in mammalian heart and liver. Genomics 2007, 89:70–80.

104. Font-Burgada J, Shalapour S, Ramaswamy S, Hsueh B, Rossell D, Umemura A, Taniguchi K, Nakagawa H, Valasek MA, Ye L, et al: Hybrid Periportal Hepatocytes Regenerate the Injured Liver without Giving Rise to Cancer. Cell 2015, 162:766–779.

105. Lin S, Nascimento EM, Gajera CR, Chen L, Neuhofer P, Garbuzov A, Wang S, Artandi SE: Distributed hepatocytes expressing telomerase repopulate the liver in homeostasis and injury. Nature 2018, 556:244–248.

106. Haghverdi L, Buettner F, Theis FJ: Diffusion maps for high-dimensional single-cell analysis of differentiation data. Bioinformatics 2015, 31:2989–2998.

107. Saelens W, Cannoodt R, Todorov H, Saeys Y: A comparison of single-cell trajectory inference methods. Nat Biotechnol 2019, 37:547–554.

108. Tanami S, Ben-Moshe S, Elkayam A, Mayo A, Bahar Halpern K, Itzkovitz S: Dynamic zonation of liver polyploidy. Cell Tissue Res 2017, 368:405–410.

109. Wolf FA, Hamey FK, Plass M, Solana J, Dahlin JS, Gottgens B, Rajewsky N, Simon L, Theis FJ: PAGA: graph abstraction reconciles clustering with trajectory inference through a topology preserving map of single cells. Genome Biol 2019, 20:59.

110. Lafzi A, Moutinho C, Picelli S, Heyn H: Tutorial: guidelines for the experimental design of single-cell RNA sequencing studies. Nat Protoc 2018, 13:2742–2757.

111. Lacar B, Linker SB, Jaeger BN, Krishnaswami SR, Barron JJ, Kelder MJE, Parylak SL, Paquola ACM, Venepally P, Novotny M, et al: Nuclear RNA-seq of single neurons reveals molecular signatures of activation. Nat Commun 2016, 7:11022.

112. Miyaoka Y, Miyajima A: To divide or not to divide: revisiting liver regeneration. Cell Div 2013, 8:8.

113. Knouse KA, Wu J, Whittaker CA, Amon A: Single cell sequencing reveals low levels of aneuploidy across mammalian tissues. Proc Natl Acad Sci U S A 2014, 111:13409–13414.

114. Knouse KA, Lopez KE, Bachofner M, Amon A: Chromosome Segregation Fidelity in Epithelia Requires Tissue Architecture. Cell 2018, 175:200–211 e213.

115. Margall-Ducos G, Celton-Morizur S, Couton D, Bregerie O, Desdouets C: Liver tetraploidization is controlled by a new process of incomplete cytokinesis. J Cell Sci 2007, 120:3633–3639.

116. Celton-Morizur S, Merlen G, Couton D, Margall-Ducos G, Desdouets C: The insulin/Akt pathway controls a specific cell division program that leads to generation of binucleated tetraploid liver cells in rodents. J Clin Invest 2009, 119:1880–1887.

117. Rodrigues OR, Monard S: A rapid method to verify single-cell deposition setup for cell sorters. Cytometry A 2016, 89:594–600.

118. Gasch AP, Yu FB, Hose J, Escalante LE, Place M, Bacher R, Kanbar J, Ciobanu D, Sandor L, Grigoriev IV, et al: Single-cell RNA sequencing reveals intrinsic and extrinsic regulatory heterogeneity in yeast responding to stress. PLoS Biol 2017, 15:e2004050.

119. Mora-Castilla S, To C, Vaezeslami S, Morey R, Srinivasan S, Dumdie JN, Cook-Andersen H, Jenkins J, Laurent LC: Miniaturization Technologies for Efficient Single-Cell Library Preparation for Next-Generation Sequencing. J Lab Autom 2016, 21:557–567.

120. Lee H, Pine PS, McDaniel J, Salit M, Oliver B: External RNA Controls Consortium Beta Version Update. J Genomics 2016, 4:19–22.

121. Liao Y, Smyth GK, Shi W: featureCounts: an efficient general purpose program for assigning sequence reads to genomic features. Bioinformatics 2014, 30:923–930.

122. Johnson WE, Li C, Rabinovic A: Adjusting batch effects in microarray expression data using empirical Bayes methods. Biostatistics 2007, 8:118–127.

123. McInnes L., J. H, N. S, L. G: UMAP: Uniform Manifold Approximation and Projection. Journal of Open Source Software 2018, 3.

124. Blondel VD, Guillaume J-L, Lambiotte R, Lefebvre E: Fast unfolding of communities in large networks. Journal of Statistical Mechanics: Theory and Experiment 2008, 2008:P10008.

125. Lun AT, McCarthy DJ, Marioni JC: A step-by-step workflow for low-level analysis of single-cell RNA-seq data with Bioconductor. F1000Res 2016, 5:2122.

126. Canchola J.A., Tang S., Hemyari P., Paxinos E., E. M: Correct use of percent coefficient of variation (%CV) formula for log-transformed data. MOJ Proteomics Bioinform 2017, 6:316–317.

127. Tabula Muris C, Overall c, Logistical c, Organ c, processing, Library p, sequencing, Computational data a, Cell type a, Writing g, et al: Single-cell transcriptomics of 20 mouse organs creates a Tabula Muris. Nature 2018, 562:367–372.

